# Caregiver genotype influences larval body size in ants by modulating ecdysteroid production

**DOI:** 10.64898/2026.11.09.765415

**Authors:** Patrick K. Piekarski, Yukina Chiba, Jina Lee, Stephany Valdés-Rodríguez, Thomas S. Carroll, Daniel J.C. Kronauer

## Abstract

Parental genes shape the offspring’s rearing environment and, consequently, offspring phenotypes. However, the molecular mechanisms underlying these indirect genetic effects remain largely unknown. Here, using the clonal raider ant *Ooceraea biroi*, we uncover a neurohormonal mechanism by which caregiver genotypes affect larval body size and caste fate. Larval transcriptomics revealed caregiver-genotype effects on neuropeptides and an ecdysteroid biosynthesis gene, while enrichment analyses implicated changes in lipid metabolism. We found that the receptors of these candidate neuropeptides are expressed in the prothoracic gland, the site of ecdysteroid synthesis, and pharmacological manipulations revealed a neuropeptide-ecdysone axis that regulates body size in ants. The same hormonal axis also regulates multiple lipid-binding carrier protein genes expressed in larval fat-body cells. Together, we identify a neurohormonal pathway that is modulated by caregivers in developing larvae, linking lipid metabolism, systemic growth, and caste development.

## Introduction

The extent to which nature (i.e., genes) and nurture (i.e., environment) shape an individual’s phenotype is a fundamental biological question. However, the distinction between nature and nurture is blurred in part because an individual’s environment includes the genes of its social partners. Indirect genetic effects (IGEs) arise when the genotype of one individual affects the phenotype of another, often through social interactions (*1, 2*). This especially occurs between parents and offspring, where genes expressed in one profoundly shape the physiology and behavior of the other (*3–8*). For example, the parental genotype often affects offspring traits by shaping the rearing environment, a phenomenon known as ‘genetic nurture’ (*6*).

In many systems, parental IGEs can be quantified, and, in humans, they help explain variation in offspring educational attainment, inattention and hyperactivity, HDL cholesterol levels, height, and body mass index (*6, 9–11*). Although genetic studies identify correlations between nontransmitted parental alleles and offspring phenotypes, correlations can be confounded and causal mechanisms are rarely established (*5, 12, 13*). Cross-fostering paradigms are powerful because they enable direct tests of postembryonic, socially mediated IGEs that are distinct from germline-mediated epigenetic or prenatal influences. However, despite genomic advances and the ubiquity of genetic nurture across animals, characterizations of its molecular basis and that of IGEs more broadly are scarce and incomplete (*7, 13–16*).

The clonal raider ant, *Ooceraea biroi*, offers a tractable model for dissecting the molecular mechanisms of genetic nurture. Due to their clonal reproduction and phasic colony cycles, each generation yields cohorts of near-isogenic, age-matched individuals that are easily propagated and maintained under standardized conditions (*17*). This permits exceptional experimental control over age, genotype, and rearing environment in a highly social animal. Notably, clonal genotypes can be cross fostered to disentangle direct and indirect genetic effects (*18–20*).

Here we focus on IGEs on larvae by caregivers of two distinct but closely related clonal genotypes, A and B, which indiscriminately adopt larvae of the other genotype (*19, 20*). Larvae reared by genotype A caregivers attain larger final body sizes and are more likely to develop queen-associated morphology (*18, 19, 21*). Although queens are absent in *O. biroi*, some individuals, known as intercastes, are larger and exhibit queen-associated traits. Unlike regular workers, which have two ovarioles, intercastes possess four or more ovarioles and typically have vestigial eyes. Because body size and other caste-associated morphological and anatomical traits, such as the extent of ovary, eye and wing development, are developmentally coupled in ants (*21–25*), studying the molecular regulation of systemic growth also elucidates mechanisms of caste determination.

We uncover a neurohormonal mechanism by which caregivers influence the body size of their young, providing insight into the neuropeptidergic and endocrine control of systemic growth and caste development in ant larvae. Because many neuropeptides are ancient, pleiotropic regulators of feeding, growth and reproduction, these findings contribute to identifying molecular pathways through which parents shape offspring phenotypes across animals.

## Results

### Caregiver genotype affects offspring size and caste with few differentially expressed genes

Larvae reared by genotype A caregivers attained larger final sizes (**Fig. 1A**) and were more likely to develop as intercastes (**Fig. 1B**) despite no detectable difference in developmental duration (**Fig. 1C**). Such an effect on body size without an effect on developmental timing suggests that caregiver genotype affected larval growth rate. To test whether caregiver genotypes differ in the amount of food they provision to larvae, we quantified larval food intake by labeling food with Alexa 647, a membrane-impermeable dye that remains in the larval gut indefinitely when ingested (**Fig. 1D**). Larvae reared by different caregiver genotypes showed no detectable differences in food ingestion (**Fig. 1E**), suggesting that caregiver genotype influences larval growth through other mechanisms. Also, larval survival rates to prepupation did not significantly differ between caregiver genotypes (binomial GLM, *n* = 12 colonies per genotype; LRT: χ²(1) = 0.13, *p* = 0.718), implying that larval cannibalism did not underlie the differences in growth rate (*18–20*). Caregiver genotype therefore influenced larval size and caste development through mechanisms other than food consumption.

**Fig. 1.**
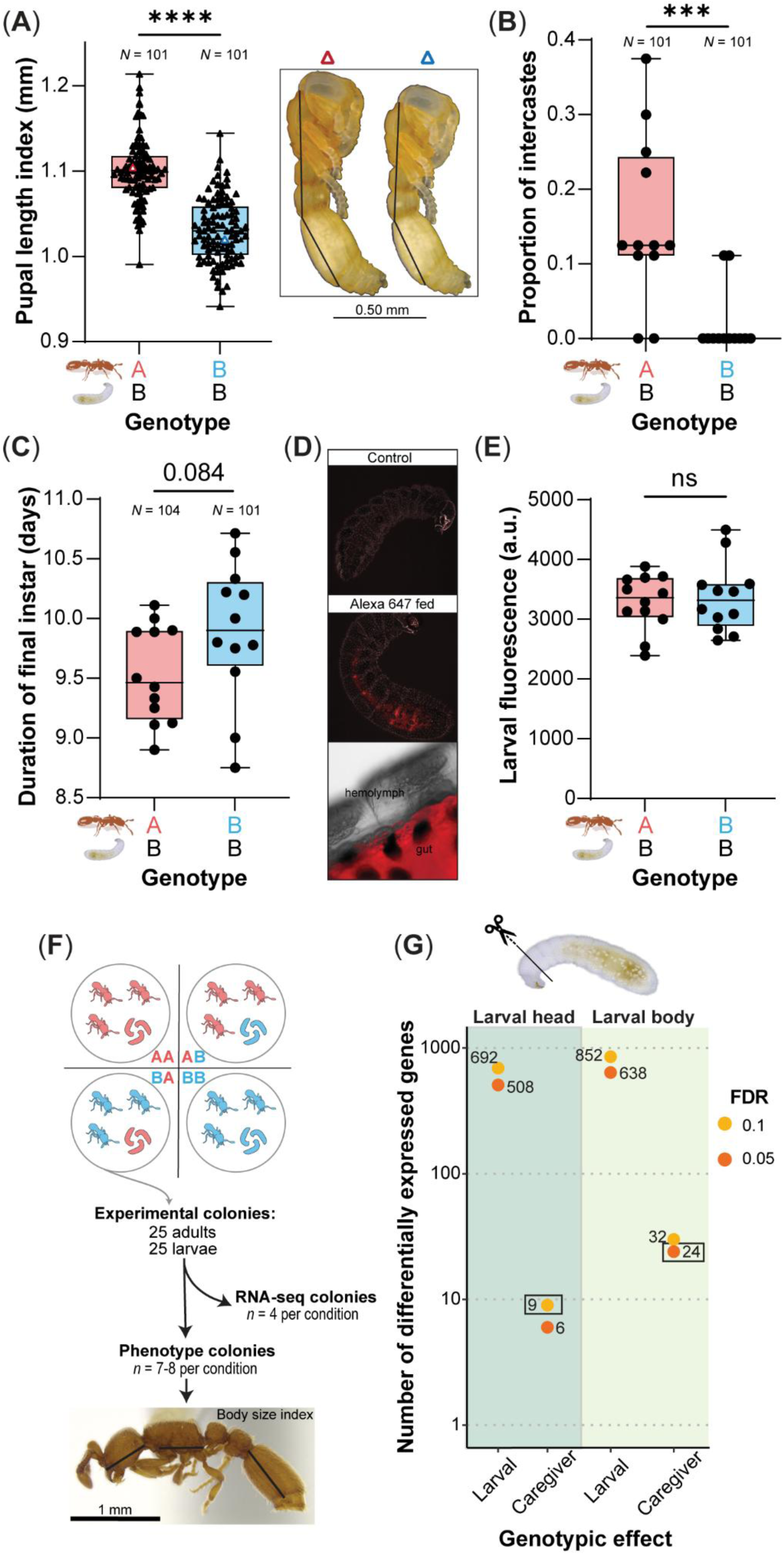
Caregiver genotype affects larval size, caste and gene expression without significantly altering feeding. (**A**) Pupal size attained by larvae reared by genotype A or B caregivers differed (LMM: χ²(1) = 48.44, *p* < 0.0001). Triangles represent individual pupae from replicate colonies (*n* = 12 per genotype). Representative pupae are shown; black lines indicate measurements summed to calculate pupal length index. (**B**) Genotype A caregivers produced a higher proportion of intercastes (binomial GLMM: χ²(1) = 12.87, *p* < 0.001; *n* = 12). (**C**) Caregiver genotype had no significant effect on larval developmental duration (LMM: χ²(1) = 2.98, *p* = 0.084; *n* = 12). (**D**) Confocal image comparing control larvae fed unlabeled food and larvae fed food infused with Alexa Fluor 647. The fluorophore is confined to the larval gut. (**E**) Relative fluorescence of larvae fed dyed (Alexa Fluor 647) food was not significantly influenced by caregiver genotype (unpaired *t*-test: *t*(22) = 0.38, *p* = 0.707). Each point represents a colony (*n* = 12). (**F**) Cross-fostering RNA-seq experimental design using clonal genotypes A and B. Ten fourth-instar larvae from a subset of colonies (*n* = 4 per condition) were collected for RNA extraction 12–24 hours after ≥50% reached the fourth instar. In the remaining ‘phenotype colonies’, focal brood were measured for adult body size, developmental duration, and survival (*n* = 7–8 colonies per condition); black lines indicate measurements summed to calculate body size index. (**G**) Number of differentially expressed genes in larval heads and bodies due to larval or caregiver genotype. In heads, 508 genes were differentially expressed due to larval genotype (P_FDR_ < 0.05), whereas only 9 genes were influenced by caregiver genotype (P_FDR_ < 0.1). In bodies, 638 and 24 genes differed by larval and caregiver genotype, respectively (P_FDR_ < 0.05). Black boxes highlight genes differentially expressed due to indirect genetic effects. Asterisks indicate significant differences (\*\*\**p* < 0.001; \*\*\*\**p* < 0.0001). All colonies contained genotype B brood.

To discover molecular factors influenced by caregiver IGEs that regulate larval growth, we conducted a cross-fostering experiment and performed RNA-seq separately on heads and bodies of early fourth-instar larvae (**Fig. 1F**). Prior to RNA-seq, we analyzed phenotypic outcomes of brood from experimental colonies set in parallel and confirmed the significant differences in final body size and intercaste proportions due to caregiver genotype (**fig. S1, A** to **D**).

RNA-seq generated ∼35.6 million mapped reads per sample (range: 31.3–48.0 million). PCA separated samples by tissue (PC1) and larval genotype (PC2; **fig. S1E**). Larvae showed differential expression due to caregiver genotype in only nine genes in the head (P_FDR_ ≤ 0.1) and 24 genes in the body (P_FDR_ ≤ 0.05) (**Fig. 1G**). The median effect sizes of IGEs were similar to or greater than those of direct genetic effects (DGEs) (**fig. S1F**). Overall, we observed a modest number of differentially expressed genes despite robust IGEs on body size, providing a focused window into the molecular factors regulating size in ants.

### Caregiver genotype alters expression of neuropeptides that regulate ecdysone synthesis in larvae

Among the genes most significantly influenced by IGEs in the larval head was *shadow* (*sad*), a Halloween gene expressed in the prothoracic gland (PG) (**Fig. 2A**). *Sad* encodes the enzyme converting 2-deoxyecdysone into ecdysone (*26*). After secretion from the PG into the hemolymph, ecdysone is converted into its active form, 20-hydroxyecdysone (20E), in peripheral tissues (*27*). Hereafter, we use the term ‘ecdysone’ to refer to both 20E and its precursor. While surges of ecdysone induce developmental transitions (*28*), basal levels of ecdysone inhibit systemic growth during larval development by repressing d*Myc*, a transcription factor and master regulator of cell metabolism and growth, and by antagonizing insulin/insulin-like signaling (IIS) in the fat body of *Drosophila melanogaster* (*29–31*). Across insects, ecdysteroid biosynthesis in the PG is largely regulated by neuropeptides, which modulate the transcription of Halloween genes (*32, 33*). In *D. melanogaster* larvae, ecdysone synthesis and release from the PG is modulated by insulin-like peptides (ILPs) (*29, 34–37*), neuropeptide F (NPF) (*38*), and prothoracicotropic hormone (PTTH) (*39–41*). Furthermore, the neuropeptides allatostatin A (AstA), corazonin (Crz), and pyrokinin-2 (PK2, encoded by *hugin*/*pban*) influence ecdysone synthesis indirectly by modulating PTTH neuron activity (*42–44*). Likewise, both *ptth* and *pban*-encoded peptides stimulate ecdysteroid synthesis in Lepidoptera (*33, 40, 45–47*), while Crz promotes *sad* expression and ecdysteroid production in the crab *Portunus trituberculatus* (*48, 49*). Remarkably, *ptth*, *crz*, and *pban* are included in the list of nine genes differentially expressed in larval heads due to caregiver genotype (P_FDR_ ≤ 0.1; **Fig. 2A, 2B**, and **fig. S2A**). Finally, *ilp2*, which encodes the canonical insulin-like peptide in ants (*50*), showed significant differential expression before multiple comparisons corrections (**Fig. 2B** and **fig. S2A**).

**Fig. 2.**
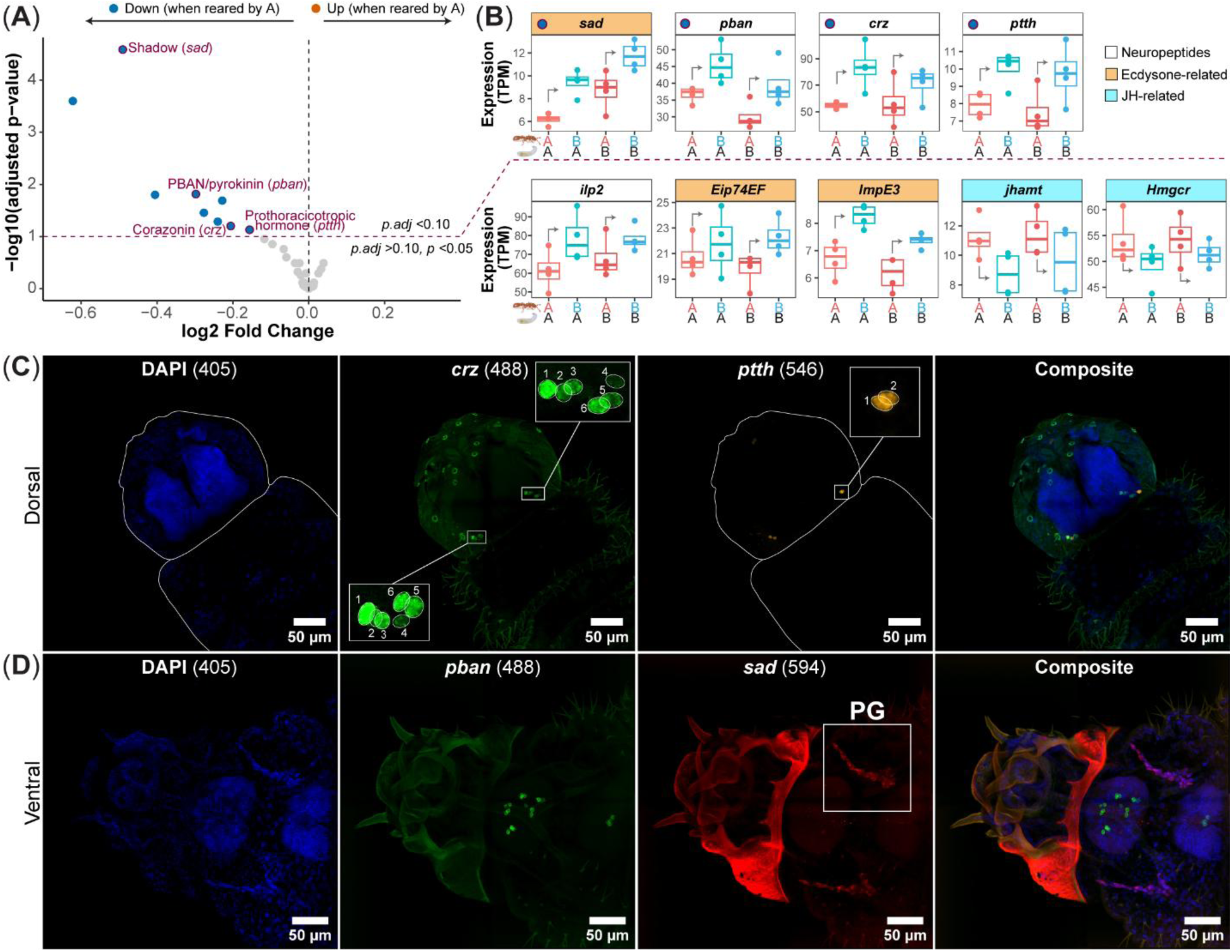
Caregiver genotype alters expression of neuropeptides that regulate ecdysone synthesis in larvae. (**A**) Volcano plot of RNA-seq from heads of early fourth-instar larvae reared by genotype A or B caregivers. The nine differentially expressed genes due to caregiver genotype (P_FDR_ < 0.1) are highlighted in blue, with genes of particular interest labeled. (**B**) Boxplots showing Salmon-estimated transcripts per million (TPM) for genes of interest across genotypic conditions. The top row shows differentially expressed genes due to caregiver genotype (P_FDR_ < 0.1), and the bottom row shows selected genes exhibiting differential expression before Benjamini-Hochberg correction in the DESeq2 analysis (*p* < 0.05, but P_FDR_ > 0.1). On the x-axis, letters denote caregiver genotype (top row) and larval genotype (bottom row). Grey arrows indicate expression differences between larvae reared by different caregiver genotypes. (**C**, **D**) RNA-FISH confocal images showing (**C**) *crz* and *ptth* expression (dorsal view), and (**D**) *pban* and *sad* expression (ventral view) in heads of early fourth-instar larvae. Nuclei were counterstained with DAPI (shown in blue). The white box indicates the prothoracic gland (PG) location. Numbers in parentheses indicate laser excitation wavelengths.

*Ilp2*, *crz*, *ptth*, and *pban* expression were significantly positively correlated across samples, but expression of *sad* was not significantly correlated with any neuropeptide (**fig. S2B**). The reason is that *crz*, *ptth*, and *pban* expression were significantly anticorrelated with *sad* expression when controlling for genotypic effects (**fig. S2, D** to **F**). Concurrently, caregiver genotype produced coordinated changes in expression of *sad*, *crz*, *ptth*, *pban*, and *ilp2*, with all genes expressed at higher levels in larvae reared by genotype B caregivers (**Fig. 2B** and **fig. S2, D** to **G**). Together, these patterns are consistent with caregiver genotype influencing ecdysteroid synthesis by altering baseline expression of *crz*, *ptth*, *pban*, and *ilp2* in larvae, with ecdysone exerting negative feedback on neuropeptide expression within conditions.

Larvae reared by genotype B caregivers exhibited elevated expression of *ecdysone-induced protein 74EF* (*Eip74EF*; LOC105284568) and *ecdysone-inducible gene E3* (*ImpE3*; LOC105277032), whereas expression of two key genes encoding juvenile hormone (JH) biosynthetic enzymes, *jhamt* (LOC105287635) and *Hmgcr* (LOC105283012), were reduced (P_FDR_ ≥ 0.1, but *p* < 0.05; **Fig. 2B**). Furthermore, *ptth*, *crz*, *pban* and *ilp2* expression were significantly anti-correlated with *jhamt* expression (**fig. S2B**). We validated the RNA-seq results by RT-qPCR of *sad, crz, jhamt,* and *ilp2* using the same RNA samples. All four genes were differentially expressed due to caregiver genotype (unpaired t-tests, one-tail: all *p* < 0.05; **fig. S2C**), confirming the RNA-seq results and corroborating that *ilp2* and *jhamt* expression were also affected by caregiver IGEs. Overall, our transcriptomic results suggest coordinated but opposing changes in expression for JH- and ecdysone-related genes due to caregiver genotype, with ecdysteroid production being partly under control by Crz, PTTH, PBAN-peptides and ILP2 (**Fig. 2B**). These findings suggest that, in the clonal raider ant, conserved neurohormonal axes regulating ecdysteroid production across insects are modulated by caregivers to regulate larval body size.

Using RNA Fluorescence In Situ Hybridization (RNA-FISH), we found that *O. biroi* larvae express *crz* in six bilateral pairs of dorsolateral neurons in the brain, but not elsewhere in the head (**Fig. 2C**). Across insects, a small, conserved set of dorsolateral (pars lateralis) protocerebral neurosecretory cells release Crz into the hemolymph from axon terminals in the corpora cardiaca (*51*). Two bilateral dorsolateral neurosecretory cells express *ptth* in ant larvae (**Fig. 2C**), as in *D. melanogaster* and Lepidoptera (*39*). Two additional cells located anteriorly in the head showed a weaker signal for *ptth* expression (**Fig. 2C**). Expression of *pban* in the larval head was limited to eight pairs of bilateral neurons (16 total) in the subesophageal ganglion and four neurons in the prothoracic ganglion of the ventral nerve cord (**Fig. 2D**). Because ants have two insulin-like peptides, *ilp1* and *ilp2* (*50*), and IIS stimulates growth, we localized expression of both genes in the larval head. *Ilp2* was expressed in 20–24 medial neurosecretory cells bilaterally, defining the canonical insulin-producing cell (IPC) population (*52*), whereas *ilp1* was expressed in a separate anterodorsal neuronal population (**Movie S1**) and several cells ventral to the IPCs (**Movie S2**). Lastly, expression of *sad* was detected exclusively in PG cells (**Fig. 2D**). To our knowledge, this study provides the first RNA-FISH-based localization of these neuropeptides in ant larvae, revealing expression patterns consistent with those previously described in other insect larvae and highlighting the conserved nature of neuroendocrine systems regulating growth and development.

### Neuropeptide receptors are expressed in the prothoracic gland

Based on these results, we conducted RNA-FISH to assess whether PG cells express the receptors of neuropeptides encoded by *crz*, *ptth*, *pban*, and *ilp2*. The genome of *O. biroi* contains a single receptor for corazonin (*crzR*; LOC105282512) and PTTH (*torso*; LOC105276519), but two receptors for the pyrokinin/PBAN family of peptides (*pk1r*/*pbanr1* and *pk2r*/*pbanr2*; LOC105283619 and LOC105283116, respectively) and insulin-like peptides (*InR1* and *InR2*; LOC105284680 and LOC105285047, respectively). Phylogenetically, InR2 of Hymenoptera corresponds to the ancestral insect insulin receptor (cluster I) and to the single remaining InR in *D. melanogaster*, whereas InR1 belongs to InR cluster II, a paralogous lineage that arose in an early pterygote ancestor (*53*). Receptors for pyrokinin/PBAN peptides belong to two groups: pyrokinin-1/diapause hormone receptor (PK1/DHr) and pyrokinin-2*/*PBAN receptor (PK2/PBANr) (*54*). ObPBANr1 belongs to the PK1/DHr receptor group, while ObPBANr2 is homologous to PK2/PBANr (**Supplementary Note 1**).

We found that cells of the ant PG, marked by *sad* expression, co-express *crzR*, *torso*, and *pbanr2*, but not *pbanr1* (**Fig. 3, A** to **D**). *InR1* was also expressed by PG cells, with broad, ubiquitous expression throughout the larval head, while *InR2* expression was not detected in any cells (**fig. S3A**). Expression of *crzR*, *torso*, *pbanr1*, and *pbanr2* was also detected in other larval tissues (**fig. S3, B** to **E**). Thus, the ant PG expresses receptors for neuropeptides encoded by *crz, ptth, pban,* and *ilp2*, consistent with them acting on the PG to stimulate ecdysone synthesis and/or release, thereby affecting body size. This provides the first demonstration that these neuropeptide receptors are expressed in the PG of a hymenopteran, supporting the idea that their regulation of ecdysteroid synthesis in the PG is ancestral and broadly conserved across insects.

**Fig. 3.**
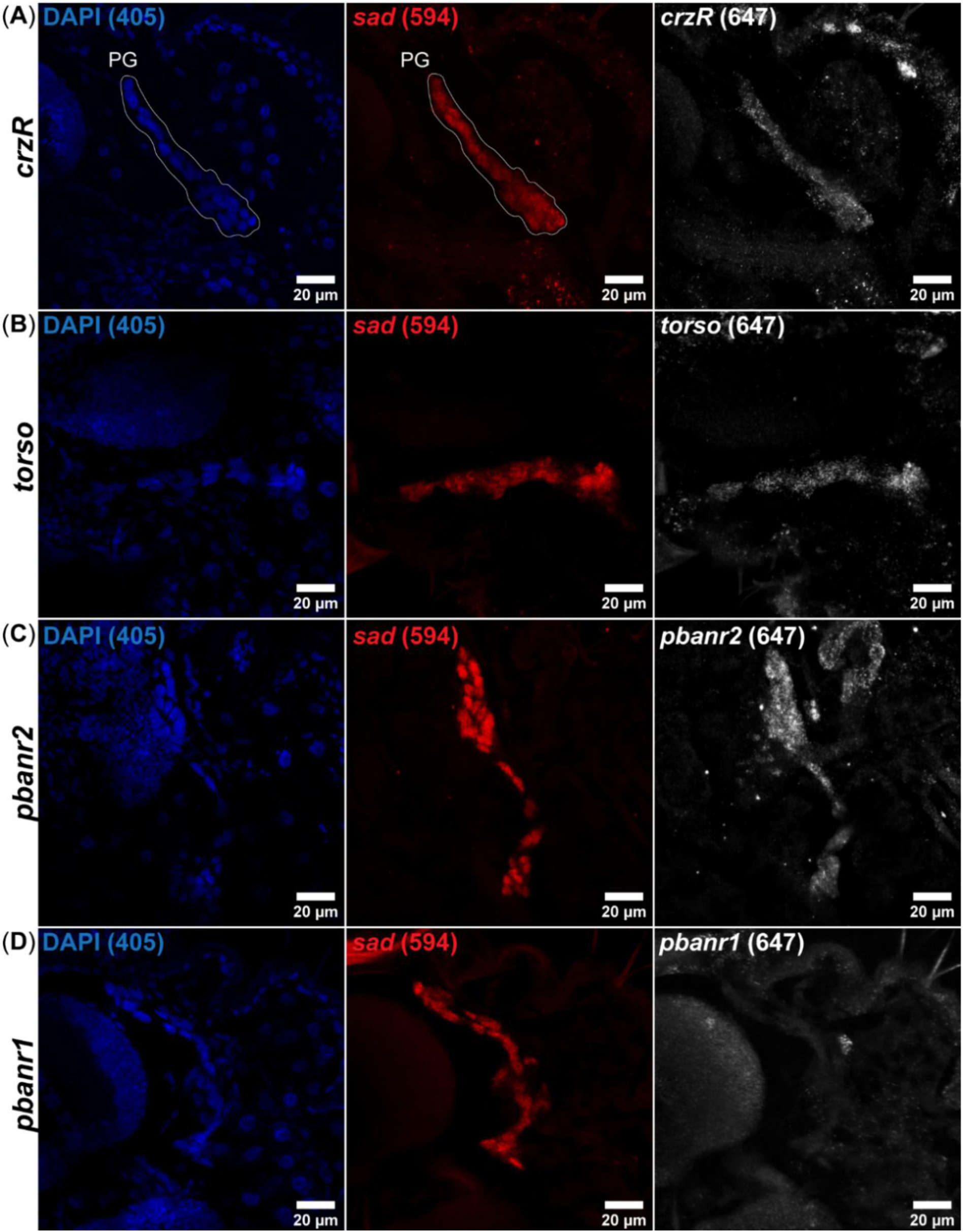
The prothoracic gland expresses receptors for Crz, PTTH, and PBAN-encoded peptides. (**A–D**) RNA-FISH confocal images of early fourth-instar larvae. Prothoracic gland (PG) cells were identified by *sad* expression (red; 594 nm), and nuclei were counterstained with DAPI (blue; 405 nm). Receptor expression is shown individually in the right column (white; 647 nm). (**A**) *crzR*, (**B**) *torso*, and (**C**) *pbanr2* were expressed in PG cells, whereas (**D**) *pbanr1* expression was not detected.

### Caregiver genotype modulates lipid-binding carrier proteins

Of the 24 genes influenced by IGEs in the larval body (P_FDR_ ≤ 0.1; **Fig. 2B**), three encode chemosensory proteins (CSPs) and three encode odorant-binding proteins (OBPs) (**Fig. 4A** and **fig. S4A**). The *O. biroi* genome contains only 15 CSPs and 15 OBPs (*55*), and both protein families are significantly overrepresented in the list of differentially expressed genes (**fig. S4a**). All differentially expressed *csp* and *obp* genes, as well as a JHBP/Takeout family gene (*jhbp*; LOC105286659), exhibited significantly higher expression in larvae reared by genotype A caregivers (**Fig. 4B**). Weighted gene co-expression network analysis (WGCNA) of larval body samples identified 31 modules, only one of which was differentially expressed across caregiver genotypes (**fig. S4B**). This module contained 39 genes, including 16 of the 24 genes recovered by differential expression analysis (**fig. S4B**). All *csp*, *obp*, and *jhbp* genes in this module show strong co-expression, as evidenced by their grouping within the same WGCNA module, suggesting a shared regulatory mechanism.

**Fig. 4.**
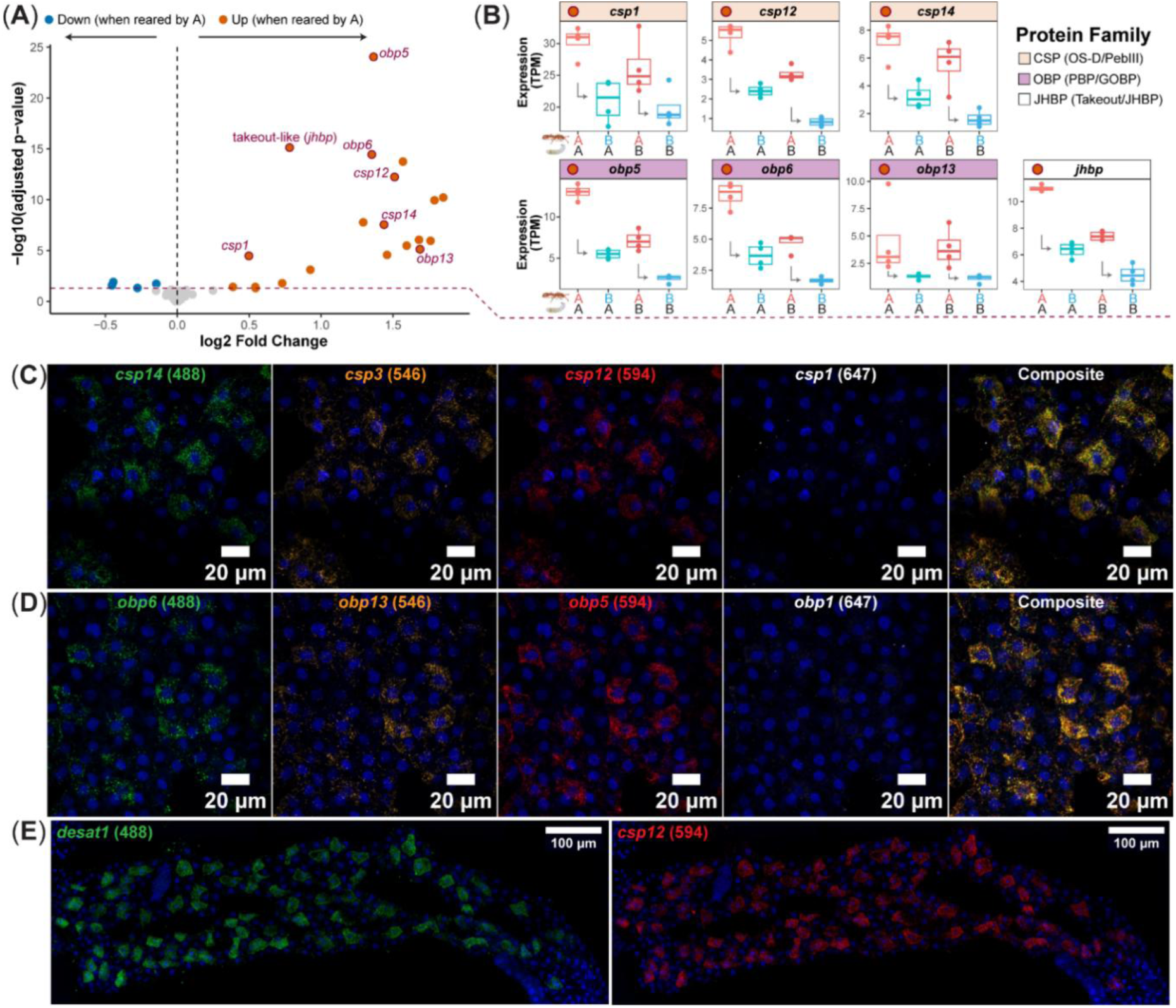
Caregiver genotype modulates expression of lipid-binding carrier protein genes in the larval body. (**A**) Volcano plot of RNA-seq from bodies of early fourth-instar larvae reared by genotype A or B caregivers. The 24 genes differentially expressed due to caregiver genotype (P_FDR_ < 0.05) are highlighted, with genes encoding secreted lipid-binding carrier proteins indicated. (**B**) Boxplots showing Salmon-estimated transcripts per million (TPM) for genes of interest across genotypic conditions. Chemosensory protein (CSP), odorant-binding protein (OBP), and takeout/juvenile hormone-binding protein (JHBP) genes are indicated in peach, purple, and white, respectively. Letters on the x-axis denote caregiver genotype (top row) and larval genotype (bottom row). Grey arrows indicate decreased expression in larvae reared by genotype B versus A caregivers. (**C–E**) RNA-FISH confocal images of whole fat bodies from early fourth-instar larvae. (**C**) Fat body cells co-express *csp14* (green), *csp3* (orange), and *csp12* (red), whereas *csp1* (white) was not detected. (**D**) *Obp6* (green), *obp13* (orange), and *obp5* (red) are co-expressed, while *obp1* (white) was not detected. (**E**) A subpopulation of fat body cells co-expresses *desat1* (LOC105283112; green) and *csp12* (red). Nuclei were counterstained with DAPI (405 nm; blue). Numbers in brackets indicate excitation wavelengths.

CSPs and OBPs are lipid-binding carrier proteins that transport pheromones and other odorants, hormones, fatty acids, and nutrients (*56*). They are expressed in a variety of adult and larval tissues in ants and other insects, including chemosensory organs, the gut, fat body, and glands (*55–57*). Although they canonically play a role in olfaction, they also function in immunity, reproduction, development, and nutrient transport (*56, 58, 59*), and in honeybees and ants, in embryonic integument formation and molting (*60, 61*). Proteins of the JHBP/Takeout superfamily are likewise secreted proteins that bind lipophilic compounds (*62–64*), and are implicated in coordinating feeding behavior and metabolism with circadian rhythm (*63, 65*).

To identify tissues in the larval body expressing the CSP and OBP genes of interest, we performed RNA-FISH. We found that the larval fat body expresses *csp3*, *csp12*, *csp14*, *obp5*, *obp6*, and *obp13*, but not *csp1* or *obp1* (**Fig. 4, C** and **D**). Co-staining for *obp6* and *csp12* further revealed that the *csp*- and *obp*-expressing cells constitute the same population (**fig. S4C**). In many insects, the fat body primarily consists of trophocytes and can include additional specialized cell types such as oenocytes and urocytes (*66*). Oenocytes are secretory cells that synthesize CHCs (*67*). They also exhibit hepatocyte-like functions, contributing to fatty acid metabolism and systemic lipid homeostasis by processing fatty acids mobilized from trophocyte triglyceride stores during starvation (*68–71*). Urocytes, by contrast, are specialized cells that store urate (*66*).

Cells co-expressing *csp*/*obp* genes (e.g., *csp12*) also expressed *desaturase 1* (**Fig. 4E**), an oenocyte marker in *D. melanogaster* (*69, 72*). These *desat1^+^*/*csp*^+^/*obp*^+^ cells correspond to conspicuous opaque white cells distributed throughout the larval fat body (**fig. S4, D** and **E**). In ants, these cells have been ambiguously referred to as both urocytes (*73–75*) and oenocytes (*76–78*). *Ecdysone Receptor* (*EcR*) expression was also enriched in these *desat1^+^/csp^+^/obp^+^*cells (**fig. S4F**). Desaturases contribute to lipid and CHC biosynthesis by generating monounsaturated fatty acids (*67, 79*). In *D. melanogaster* larvae, dMyc’s stimulatory effects on lipid synthesis in the fat body and systemic growth are dependent on *desat1* (*80*), and d*Myc* expression and function are repressed by EcR signaling (*30*). In *D. melanogaster* adults, oenocyte *desat1* promotes an unsaturated circulating lipidome, supporting triglyceride storage in trophocytes and enabling proper lipoprotein trafficking required for starvation-induced lipid droplet formation in oenocytes (*71*).

Because *EcR* is expressed in *desat1*^+^ cells of the fat body, differences in basal ecdysone levels due to caregiver genotype could be affecting lipid homeostasis. In support of this, gene set enrichment analysis (GSEA) identified significantly enriched GO terms related to lipid metabolism (**fig. S5**). Larvae reared by genotype A caregivers showed a transcriptional state associated with increased lipid catabolism, including fatty acid oxidation, along with elevated ribosomal and translational activity, consistent with enhanced metabolic fueling of growth (**fig. S5**). Given their expression in *desat1*^+^ cells (**Fig. 4E**), their affinity for binding fatty acids (*81–84*), and differential expression in a context where lipid metabolic processes were affected (**Fig. 5A** and **fig. S5**), we propose that CSPs and OBPs may play a role in lipid homeostasis.

**Fig. 5.**
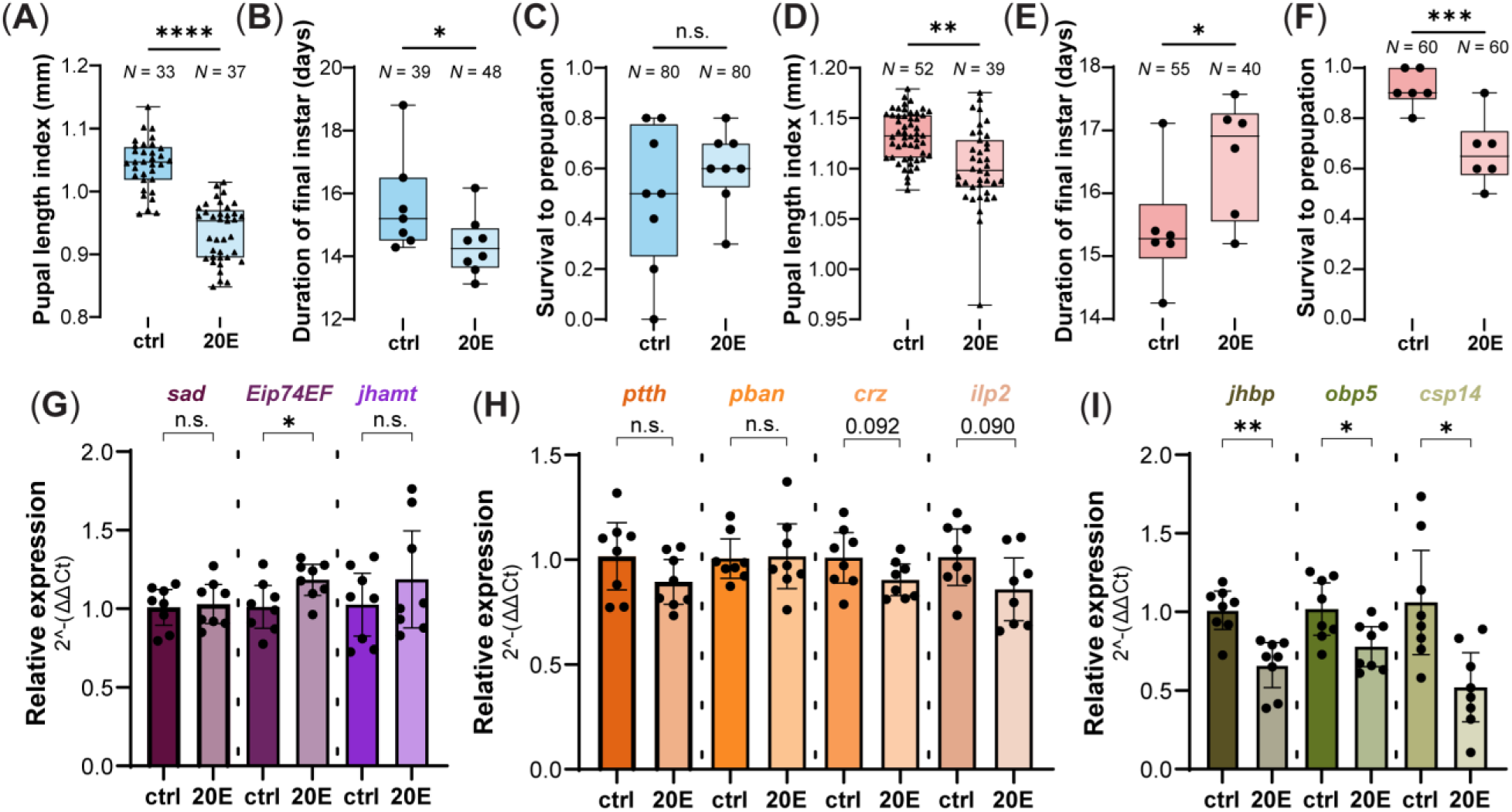
Ecdysone reduces body size and represses lipid-binding carrier protein gene expression. (**A–C**) Effects of dietary 20E (dissolved in 10% EtOH) on (**A**) body size, (**B**) developmental timing, and (**C**) survival in genotype B larvae (*n* = 8 colonies). Dietary 20E reduced pupal size (LMM: χ²(1) = 23.71, *p* < 0.0001) and decreased the duration of the final instar (CLMM: χ²(1) = 4.54, *p* = 0.033) but had no significant effect on larval survival (beta-binomial GLM: χ²(1) = 1.05, *p* = 0.306). (**D–F**) Effects of 20E on (**D**) body size, (**E**) developmental timing, and (**F**) survival in genotype A larvae (*n* = 6 colonies). Compared to solvent-only controls, 20E treatment reduced size (LMM, χ²(1) = 8.60, *p* = 0.003), prolonged development (LMM, χ²(1) = 4.60, *p* = 0.032), and reduced survival (GLM, χ²(1) = 12.02, *p* < 0.001). (**G–I**) Effects of dietary 20E on gene expression in genotype A larvae quantified by RT-qPCR (*n* = 8 colonies). (**G**) 20E treatment did not significantly affect expression of *sad* (unpaired t-test: *t*(14) = 0.29, *p_adj_* = 0.432) or *jhamt* (*t*(14) = 0.97, *p_adj_* = 0.432), whereas it increased *Eip74EF* expression (unpaired t-test, one-tailed: *t*(14) = 2.92, *p_adj_* = 0.037). (**H**) Larvae treated with 20E showed a trend toward decreased *crz* and *ilp2* expression (unpaired t-tests, one-tailed: *crz*, *t*(14) = 1.70, *p_adj_* = 0.092; *ilp2*, *t*(14) = 1.82, *p_adj_* = 0.090), whereas *ptth* and *pban* expression did not differ between treatments (unpaired t-tests, one-tailed: *pban*, *t*(14) = 0.03, *p_adj_* = 0.488; *ptth*, *t*(14) = 1.43, *p_adj_* = 0.125). (**I**) 20E treatment significantly decreased *jhbp*, *obp5*, and *csp14* expression (unpaired t-tests, one-tailed: *jhbp*, *t*(14) = 3.78, *p_adj_* = 0.010; *obp5*, *t*(14) = 2.68, *p_adj_* = 0.030; *csp14*, *t*(14) = 2.99, *p_adj_* = 0.025). P-values were corrected using the Benjamini–Hochberg FDR method. For qPCR data, bars represent the mean ± 95% CI. Asterisks indicate significant differences (\**p* < 0.05; \*\**p* < 0.01; \*\*\**p* < 0.001; \*\*\*\**p* < 0.0001).

### Ecdysone reduces body size and represses lipid-binding carrier protein gene expression

Perturbations of ecdysone synthesis or signaling alter final body size in *D. melanogaster* by changing the duration of larval growth through effects on developmental timing, altering growth rate, or both (*29, 34–37, 39–41, 44, 85–87*). To determine ecdysone’s effects on final body size and developmental timing in *O. biroi*, we fed early fourth-instar larvae of genotype B five times (once every 48 hours) with food infused with 20E or solvent control. Compared to the solvent control, 20E-treated larvae attained significantly smaller final body sizes on average (**Fig. 5A**) and had modestly accelerated development (**Fig. 5B**), without detectable effects on survival (**Fig. 5C**). For genotype A larvae, three 20E feedings significantly reduced average body size relative to controls (**fig. S6A**), without significantly affecting developmental timing or survival (**fig. S6, B** and **C**).

To assess whether caregiver-genotype effects on ecdysteroid production in larvae could explain differences in *csp* and *obp* expression, and whether candidate neuropeptides are reciprocally regulated by ecdysone, we repeated the 20E-feeding experiment while collecting a subset of colonies for RT–qPCR. As before, 20E-fed larvae attained smaller final body sizes than controls on average (**Fig. 5D**). However, in this instance, larvae in the 20E treatment group exhibited longer development (**Fig. 5E**) and a significantly reduced survival rate (**Fig. 5F**). Because 20E-fed larvae attained smaller body sizes despite a longer developmental duration, the observed size reduction is unlikely due to a shorter duration of larval growth. Overall, dietary 20E consistently reduced body size despite variable effects on developmental timing, consistent with ecdysone reducing body size, at least in part, by inhibiting larval growth in *O. biroi*.

20E treatment significantly increased expression of the early response gene *Eip74EF* (**Fig. 5G**), indicating a cellular response to experimentally elevated 20E levels (*29, 34*). 20E-treated larvae also exhibited a trend toward decreased *crz* and *ilp2* expression (**Fig. 6H**). Lastly, dietary 20E significantly repressed *jhbp*, *obp5*, and *csp14* expression (**Fig. 6I**). This result links the transcriptional changes indicative of caregiver genotype effects on ecdysteroid synthesis (in the larval head) with changes in the expression of lipid binding carrier protein genes (in the larval body).

**Fig. 6.**
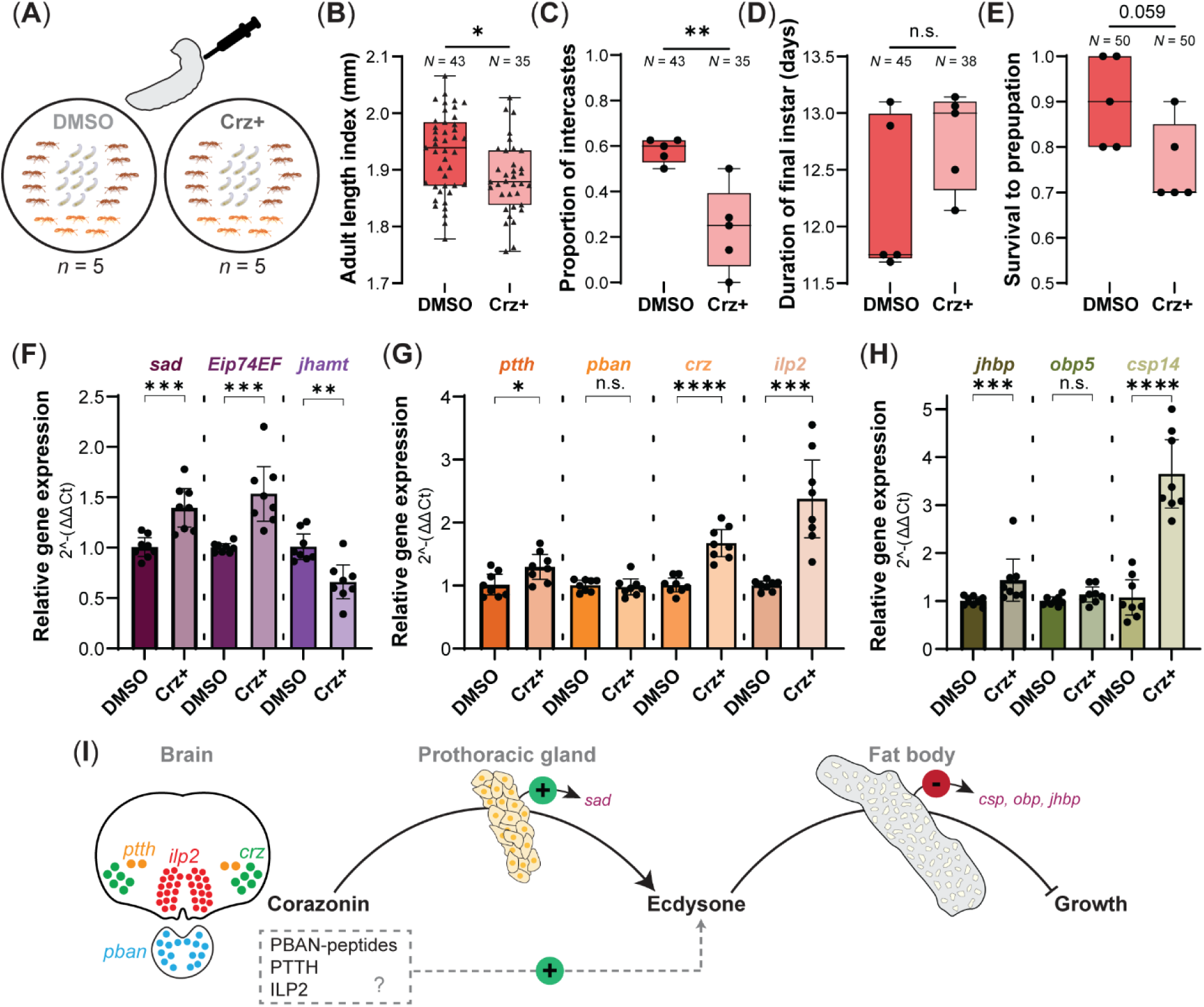
Corazonin reduces body size and upregulates *shadow*. (**A**) Experimental design for corazonin injections. Early fourth-instar larvae were injected with vehicle (0.5% DMSO in artificial hemolymph containing 1% blue food coloring; DMSO control) or ∼10 ng of corazonin in the same vehicle (Crz+). Experimental colonies (*n* = 5 per condition) comprised ten injected larvae with ten older and five callow workers (indicated by darker and lighter color, respectively). (**B–E**) Effects of corazonin injection on (**B**) adult body size, (**C**) intercaste proportion, (**D**) duration of the final instar, and (**E**) survival to prepupation. Crz injection reduced adult body size (LMM: *χ*²(1) = 4.32, *p* = 0.038) and intercaste proportions (binomial GLMM: χ²(1) = 9.59, *p* = 0.002), did not significantly affect developmental timing (LMM: *χ*²(1) = 2.34, *p* = 0.127), and showed a trend toward reduced survival (binomial GLM: *χ*²(1) = 3.56, *p* = 0.059) relative to 0.5% DMSO controls. (**F–H**) Effects of Crz injection on larval gene expression quantified by RT-qPCR (*n* = 8 colonies). (**F**) Crz treatment increased expression of *sad* (unpaired t-test, one-tailed: *t*(14) = 4.64, *p_adj_* < 0.001) and *Eip74EF* (Welch’s t-test, one-tailed: *t*(7.8) = 5.64, *p_adj_* < 0.001), and decreased *jhamt* relative to controls (*t*(14) = 3.70, *p_adj_* = 0.003). (**G**) Crz injection increased *ptth*, *crz*, and *ilp2* expression (unpaired t-tests: *ptth*, *t*(14) = 2.62, *p_adj_* = 0.025; *crz*, *t*(14) = 7.02, *p_adj_* < 0.0001; *ilp2*, Welch’s *t*(7.9) = 7.14, *p_adj_* < 0.001) but did not significantly affect *pban* expression (*t*(14) = 0.50, *p_adj_* = 0.625). (**H**) Crz significantly increased expression of *jhbp* (Mann–Whitney test: *U* = 2, *p_adj_* = 0.001) and *csp14* (unpaired t-test: *t*(14) = 7.75, *p_adj_* < 0.0001) but not *obp5* (unpaired t-test: *t*(14) = 1.72, *p_adj_* = 0.11). P-values were corrected using the Benjamini–Hochberg FDR method. Bars represent the mean ± 95% CI. Experiments were performed with genotype A ants. Asterisks indicate significant differences (\**p* < 0.05; \*\**p* < 0.01; \*\*\**p* < 0.001; \*\*\*\**p* < 0.0001). (**I**) The proposed hormonal axis, which is modulated by caregiver genotype and regulates larval size in *O. biroi*. Crz released from neuroendocrine cells acts on the prothoracic gland to promote *sad* expression and, therefore, ecdysteroid production. Ecdysone in turn inhibits growth, likely via the fat body, in part through repressing the expression of lipid-binding carrier protein genes (e.g. *csp*, *obp* and *jhbp* genes). The action of other candidate neuropeptides (PBAN, PTTH, and ILP2) remain to be verified experimentally in ants.

### Corazonin reduces body size and upregulates shadow

To test whether Crz increases *sad* expression and reduces larval body size, we injected early fourth-instar larvae once with ∼10 ng of synthetic Crz (**Fig. 6A**). These larvae attained smaller adult sizes on average compared to controls injected with solvent control (**Fig. 6B**) and had a significantly reduced probability of becoming intercastes (**Fig. 6C**). Crz treatment had no significant effect on developmental duration (**Fig. 6D**) and showed a trend towards reduced survival to prepupation (**Fig. 6E**). Differences in survival across colonies did not explain variation in the average body size of ants (R² = 0.02, *p* = 0.709).

We then repeated the Crz injection experiment to corroborate that Crz reduces body size, while also harvesting a subset of colonies for gene expression analysis using RT-qPCR. Crz treatment again resulted in smaller pupal body sizes on average compared to controls (**fig. S6D**) with no significant effect on developmental duration (**fig. S6E**). However, in this case, Crz injection significantly decreased larval survival (**fig. S6F**). Together, our results show that Crz reduces body size without detectable effects on developmental duration, suggesting that Crz inhibits growth in *O. biroi*.

Our gene expression analyses revealed that Crz upregulates *sad* in the PG, with a corresponding increase in *Eip74EF* expression (**Fig. 6F**), supporting an increase of hemolymph ecdysone levels. These analyses further revealed that Crz either directly or indirectly suppresses *jhamt* (**Fig. 6F**), and that this effect is likely independent of its impact on ecdysone levels, given that larvae fed with 20E had no detectable change in *jhamt* expression (**Fig. 5G**).

While Crz treatment had no effect on *pban* expression, it did significantly increase *ptth*, *crz*, and *ilp2* expression (**Fig. 6G**), consistent with the correlated expression of these neuropeptides in developing larvae (**fig. S2B**). In *D. melanogaster*, Crz neurons respond to nutrition, sensing glucose (*88*) and fructose (*89, 90*), and receiving upstream octopamine signals (*43*). Fat body crzR signaling modulates expression of ILPs in IPCs (*91*), and Crz/sNPF neurons directly regulate IPC and PTTH neuron activity (*43, 88, 92*). Our results show that, in *O. biroi*, *crz* and *ilp2* expression are tightly coupled in early fourth-instar larvae, but the mechanistic basis of this correlation remains unknown.

Crz treatment increased the expression of *jhbp* and *csp14*, whereas the increase in *obp5* was not statistically significant (**Fig. 6H**). Because 20E treatment suppressed *jhbp*, *csp14*, and *obp5* expression (**Fig. 5I**), these findings suggest that Crz signaling may partially counteract ecdysone’s inhibitory effects on these genes. Consistent with this, in *Drosophila*, Crz acts as a neuroendocrine regulator of metabolic homeostasis via the fat body during nutritional stress (*91*). In addition, Crz knockdown in *Drosophila* larvae and Crz neuron ablation in adults increase carbohydrate and lipid storage, respectively (*88, 91–93*). Together, these observations raise the possibility that Crz and ecdysone signaling interact to maintain metabolic homeostasis in ants.

Altogether, our results support a neurohormonal pathway affecting body size and caste development that is modulated by caregiver genotype and that involves corazonin and ecdysone. In our proposed model, Crz promotes ecdysteroid synthesis in the PG, and ecdysone in turn inhibits larval growth, likely through effects on the fat body (**Fig. 6I**).

## Discussion

We describe an instance of genetic nurture where a neurohormonal pathway that controls offspring size is modulated by caregiver genotype. This modulation occurs via effects on the expression of neuropeptides that regulate ecdysteroid synthesis across arthropods, including *crz, ptth, pban,* and *ilp2*, as well as the biosynthetic gene *sad*. We further show that receptors of these neuropeptides are expressed in the ant prothoracic gland and, through ecdysone and corazonin manipulations, identify a corazonin–ecdysone hormonal axis that regulates body size and caste development. Together, these findings suggest that caregiver IGEs affect the neuroendocrine regulation of ecdysteroid production during development, thereby altering final body size.

In *D. melanogaster*, *crz* neurons regulate basal ecdysteroid production via PTTH neurons, and inhibition of Crz signaling increases growth rate without significantly affecting pupariation timing (*43*). In the swimming crab *P. trituberculatus*, Crz directly promotes *sad* expression and increases ecdysteroid production in the Y-organ, the crustacean homolog of the PG (*48, 49*). Expression of *crzR* has also been reported in the PG of the kissing bug (*94*). Our findings that Crz regulates ecdysone synthesis (at least in part via *sad*) and body size in *O. biroi* thus suggest that Crz has an ancient role in modulating ecdysteroid synthesis.

This identifies Crz as an evolutionarily conserved neuroendocrine factor regulating metabolic homeostasis, growth and maturation across invertebrates, with parallels to its mammalian homolog gonadotropin-releasing hormone (GnRH) (*95–98*). In mammals, the onset of puberty is gated by hypothalamic GnRH neurons, with kisspeptin acting as a key upstream signal that stimulates GnRH secretion in response to metabolic cues (*99–101*). In rats, natural variation in maternal licking/grooming affects offspring pubertal timing and is accompanied by differences in GnRH neuron number and downstream hormonal activity (*102*), and maternal separation induces premature increases in kisspeptin and GnRH expression (*103*). Naked mole-rats exhibit a reproductive division of labor in which only the queen reproduces, with social inhibition of hypothalamic GnRH release in subordinates preventing the onset of puberty (*104*). It is therefore intriguing to consider whether socially mediated IGEs could influence offspring growth and pubertal timing by modulating kisspeptin–GnRH signaling in mammals, including humans. Observations across animals point to Crz/GnRH signaling as a conserved neuroendocrine pathway that regulates metabolism, growth, and reproduction (*95–98*). Here, we show that, in the clonal raider ant, this pathway can be modulated in larvae by caregiving adults.

We also identified other neuropeptides modulated by caregiver genotype, namely PTTH, PBAN, and ILP2, that may regulate ecdysteroid production and final body size in *O. biroi*. These are likely conserved roles of these ancient neuropeptides. In *Bombyx mori* and *D. melanogaster* larvae, activation of *torso* by PTTH stimulates ecdysone synthesis in the PG by upregulating expression of Halloween genes (*33, 39–41, 45, 105*). In *D. melanogaster*, loss of PTTH/Torso signaling delays larval development and typically increases final body size through prolonged feeding (*39–41*). Similarly, inhibition of PTTH signaling on the PG in hemipteran bugs reduces ecdysone titers, prolonging the nymphal stages and increasing body size (*106, 107*). In moths, diapause hormone, one of several FXPRL-amide peptides encoded by the *pban* gene, stimulates ecdysteroid production in the PG via its receptor (*46, 47*). In *Drosophila*, pyrokinin-2 is encoded by *hugin* (ortholog of *pban*) and modulates PTTH neuron activity to influence developmental duration and body size (*44, 108, 109*). Lastly, in *Drospohila* and *Bombyx* larvae, IIS/TOR signaling in the PG promotes ecdysteroid production (*29, 35–37, 85*). Because heightened IIS in the PG is associated with reduced systemic growth, this may explain why larvae reared by genotype B caregivers exhibit relatively higher *ilp2* expression yet attain smaller body size.

Perturbations of IIS in the PG can affect both body size and developmental timing in some contexts (*34, 35*), but primarily body size in others (*29, 36, 86*). Together, this indicates that PG InR can regulate both basal ecdysone levels that inhibit systemic growth and alter the timing of the ecdysone peak associated with attainment of critical weight (*29, 36, 37, 86, 87*). Similarly, our ecdysone treatments robustly reduced body size, with variable effects on developmental timing. Although we could not definitively determine whether our ecdysone manipulation affected body size through changes in basal ecdysone levels, the timing of an ecdysone peak, or both, our results collectively suggest that elevated basal ecdysone levels likely contributed, at least in part, to the observed reduction in body size.

Relative to JH, there has been limited research on ecdysone’s role in ant larval growth and caste development. To our knowledge, there is only a single functional study performed over 50 years ago (*110*). Although scarce, all studies are consistent with ecdysone suppressing growth and queen development. In *Pheidole* and *Pogonomyrmex* ants, worker-biased eggs have higher ecdysteroid levels than queen-biased eggs (*111, 112*). In *Plagiolepis pygmaea*, ecdysteroid titers during late larval development are higher in worker larvae compared to queen larvae (*113*). Lastly, injection of ecdysone into queen-destined larvae in the final instar shortens development, decreases body size, and causes some adults to emerge as workers in *Myrmica rubra* (*110*). Here we report experiments in which dietary 20E resulted in ants with smaller body sizes, consistent with ecdysone suppressing larval growth in ants.

Across ants, JH-analog treatment generally promotes larger body size and/or queen (or soldier) development (*25, 110, 114–119*). JH typically increases body size by extending larval development in ants (*25, 110, 118*) and other insects (*120, 121*), consistent with its anti-metamorphic function (*28, 122–124*). Indeed, JH and ecdysone are linked by reciprocal feedback and shared neuroendocrine regulation: JH can inhibit ecdysteroid biosynthesis and PTTH signaling, ecdysone can suppress JH synthesis, and both pathways are jointly modulated by conserved pleiotropic neuropeptides (*97, 122, 125–130*). Although the mechanisms underlying their opposing changes in ants remain unclear, taken together with previous studies, our findings support an interplay between JH and ecdysone throughout development with stage-specific outcomes. During embryonic and larval development, a high-ecdysone, low-JH state biases individuals toward smaller size and worker-like morphology, whereas in adults, a similar hormonal profile supports queen-like reproductive physiology and brain state (*25, 110, 112, 116, 131–134*). Also, in adult ants and paper wasps, Crz is upregulated in the brains of non-reproductives compared to reproductives, and Crz injection induces worker-like behavior and suppresses oocyte maturation in *Harpegnathos saltator* ants (*132, 135*). While Crz promotes worker-like behavior and physiology in adults, its role during ant larval development was previously unknown. Our discovery that Crz inhibits larval growth and decreases the probability of developing queen-associated morphology in *O. biroi* suggests that Crz promotes worker-associated phenotypes in both ant larvae and adults. Overall, our results support the idea that evolution has co-opted pleiotropic regulators of feeding, growth, and reproduction, including JH, ecdysone and corazonin, to regulate both larval caste determination and adult behavioral division of labor (*97*).

Our study also uncovered that elevated ecdysone represses *csp*/*obp* expression in ant larvae. Intriguingly, reduced ecdysone signaling induces *obp* expression during olfactory system development in *M. sexta* pupae (*136, 137*). Also, in *Drosophila* S2 cells, treatment with 20E completely suppresses expression of the CSP family gene *phk-3* (*138*). Together, this raises the question of whether CSPs and OBPs are transcriptionally regulated by ecdysone signaling across tissues and species. Furthermore, we found that multiple *csp* and *obp* genes exhibit correlated expression, are enriched in a distinct population of *desat1*^+^ cells in the fat body, and respond to caregiver IGEs and ecdysone. While their roles beyond olfaction remain largely a mystery (*56, 58*), our study implicates ant CSPs and OBPs as potentially important for fatty acid mobilization and metabolism. Notably, *D. melanogaster* Obp44a was recently shown to be a glia-derived, secreted, non-canonical fatty-acid binding protein important for brain lipid homeostasis, functioning as both an intracellular and extracellular lipid chaperone (*82*). It is further plausible that ecdysone’s effect on size is in part mediated via its repression of genes encoding lipid-binding carrier proteins in the fat body. Consistent with this, knockout of a fat-body OBP in the fall armyworm moth *Spodoptera frugiperda* reduces body size (*139*). We propose that ecdysone may repress *csp* and *obp* expression in the fat body, which reduces lipid mobilization and catabolism and decreases systemic growth.

Lastly, CSPs and OBPs are frequently associated with ant supergenes that underlie polymorphisms in social structure and caste development (*140–143*). So far, this finding has been interpreted in the context of chemical communication. However, direct experimental evidence showing that CSPs and OBPs regulate social polymorphisms by affecting chemical communication is lacking. Our findings suggest an alternative hypothesis, namely that CSPs and OBPs act during development to regulate the alternative caste phenotypes frequently associated with ant supergenes.

## Methods and Materials

### Study model, maintenance, and genotypes used

Stock colonies of *O. biroi* were maintained in climate-controlled rooms at 25 ± 1 °C and ≥60% relative humidity in sealed plastic containers with a plaster of Paris floor. Stock colonies were supplied with frozen fire ant (*Solenopsis invicta*) larvae and pupae as food three times per week during the foraging phase. Genotypes A and B were collected in Okinawa, Japan and St. Croix, U.S. Virgin Islands, respectively, but both lines belong to a clade native to Bangladesh and have successfully invaded other regions (*144, 145*). Clonal genotypes A and B differ by ∼0.38 base pairs (bp) per 1,000 bp (unpublished data), while two unrelated humans differ by ∼1 bp per 1,000 bp on average. Due to their close relationship (*17, 144, 145*), no major incompatibility occurs between the adults and larvae of the different genotypes when conducting cross-fostering experiments (*18–20*). Unless otherwise noted, experimental colonies were established in 50 mm Petri dishes with a plaster of Paris floor and maintained at 25 ± 1 °C. This study examined the effects of caregiver genotype on larval transcriptomes, growth, developmental timing, and caste fate. Although *O. biroi* lacks queens, some workers develop as larger intercastes with four or more ovarioles and vestigial eyes, while regular workers have fewer than four ovarioles and lack vestigial eyes. The larval stage of *O. biroi* comprises four instars, with most growth occurring during the final instar. Because experiments showed that caregiver effects only during the late third instar through prepupation period were sufficient to produce body size differences, all subsequent molecular profiling and functional manipulations were performed using fourth-instar *O. biroi* larvae. Potential caregiver or treatment effects specific to earlier larval stages were not evaluated.

### Differential feeding assay

Food items (*S. invicta* worker pupae) were fluorescently labeled by soaking them in 0.5 mg/mL Donkey anti-Goat IgG (H+L) Cross-Adsorbed Secondary Antibody, Alexa Fluor 647 (Invitrogen) overnight at room temperature, followed by 24 hours at 4 °C. Labeled food was stored at −20 °C in the dark. Prior to feeding, food was thawed and rinsed three times with ddH₂O.

Groups of 25 young workers (∼12 days of age) of genotypes A (colony C16) and B (colony STC6) were housed separately and fed regular, undyed food every 48 hours until eggs were laid. Once larvae hatched, feeding resumed every 48 hours. When all larvae in each colony reached the fourth instar, brood was swapped with 25 late third-instar larvae from a genotype B (STC6) stock colony. Thus, colonies (n = 12 per treatment) consisted of 25 age-matched caregivers (of genotype A or B) and 25 third-instar larvae of genotype B from the same stock. Dyed food items were administered in three feedings: 8 pupae on the day of the brood swap, 10 pupae after 48 hours, and 12 pupae after another 48 hours. Petri dishes were wrapped in aluminum foil to protect the fluorophore from light.

Twenty-four hours after the final feeding, 15 larvae were harvested per colony. The remaining individuals (8 to 10 larvae) were left in the colonies to complete the foraging phase and were later measured for final body size. The 15 larvae from each colony were pooled separately and homogenized in 200 μL ddH₂O using a TissueLyser II (QIAGEN). Fluorescence was quantified from 40 μL of supernatant loaded in triplicate into a black bottom 384-well plate. Readings were taken using a BioTek Synergy Neo2 plate reader with excitation at 600 nm, emission at 670 nm, and a gain setting of 135. A standard curve was generated from a 1:2 serial dilution series consisting of five concentrations of Alexa Fluor 647-conjugated antibody, each assayed in triplicate (linear regression: *y* = 2280*x* − 44.51, *R*² = 0.999). Mean raw fluorescence (RFU) per colony was calculated from three technical replicates. Negative controls consisting of homogenized larvae not exposed to fluorescent food exhibited background fluorescence levels ranging from 4–54 RFU, while positive larval samples exhibited RFU values ≥2000. All data collected from this experiment are deposited in Zenodo.

### Body size and developmental timing measurements

Three metrics were used to estimate body size in this study: pupal length index, adult body size index, and adult length index. The pupal length index was defined as the distance from the anterior end of the thorax to the anterior margin of the first gastral segment, plus the length of the first gastral segment (example shown in **Fig. 1A**). The adult body size index represents the sum of the length of the head, thorax, and first gastral segment measured from a profile view (example shown in **Fig. 1F**). The adult length index was measured from a dorsal view as the sum of the head, thorax, and first gastral segment lengths. Images of ants were acquired using a Leica Z16 APO microscope with a Leica DFC450 camera operated using Leica Application Suite v4.12.0 (Leica Microsystems, Switzerland). Body segment lengths were measured using Fiji (ImageJ) (*146*).

Developmental timing was measured using stage transitions. Colonies were inspected either every 12 or 24 hours, and the proportion of larvae that transitioned to the final instar and prepupal stage were recorded. For the differential feeding assay and RNA-seq cross-fostering experiment, duration of final instar represents the number of days from when ≥50% of larvae were fourth instar until ≥50% of surviving larvae reached the prepupal stage. For all other experiments, fourth-instar duration was inferred for individual larvae and represents the number of days from the start of the experiment (at which point the larvae had transitioned to the fourth instar less than 48 hours ago) until reaching the prepupal stage. Depending on the experiment, intercaste proportions represent the proportion of ants with vestigial eyes or four or more ovarioles. Vestigial eyes were used in the differential feeding experiment, whereas ovariole number was used in the cross-fostering RNA-seq and corazonin injection experiments. Larval survival rates were assessed by recording the proportion of larvae that survived until prepupation. Developmental and morphological phenotypic data collected in this study are available at Zenodo.

### Cross-fostering RNA-seq to quantify transcriptomic indirect genetic effects

Twenty-four colonies of each clonal genotype, A (C17) and B (STC6), were established in 50 mm Petri dishes. These colonies comprised 25 regular workers (∼four to seven days old) taken from stocks that were fed *S. invicta* brood. To synchronize the onset of egg-laying across genotypes, B callows were selected to be ∼two days older than those from genotype A (*18*). This minimized differences in developmental stage between larvae reared by different caregiver genotypes at the time of larval swapping. Additional backup colonies were established for both genotypes to serve as a source of replacement adults in case of mortality and to supplement larvae in experimental colonies that produced fewer than 25 larvae.

Colonies were fed 10 *S. invicta* pupae every two days until egg-laying began. Feeding was stopped simultaneously for genotypes A and B once the first eggs were observed. Colonies were maintained at 24 ± 1 °C during this reproductive phase. When larvae had hatched from over 75% of the eggs in colonies of both genotype A and B, a synchronized larval swap was performed. This results in a full factorial cross-fostering experimental design, with colonies of all four genotypic combinations of caregivers and brood being established. To control for the experience of larval swapping, larvae were swapped also within genotype. In other words, for example, colonies comprised of B adults and B larvae (i.e., BB) had their larvae swapped with another BB colony. These experimental colonies comprised 25 one-month-old caregiver workers and 25 first-instar larvae. They were housed in 50 mm Petri dishes at 27 ± 1 °C and were fed daily. For each of the four combinations of caregiver and larval genotype (i.e., AxA, AxB, BxB, BxA), there were either 11 or 12 replicate colonies.

To standardize sampling across colonies and conditions in which larvae develop at different rates, larvae were collected for RNA sequencing 12–24 hours after ≥50% of larvae reached the final instar. This is a clear, identifiable stage of larval development for which it was previously observed that there were genotype-specific differences in caregiver foraging behavior (*18*). Further, to reduce confounding effects of nutrition (i.e., differences in time since last larval feeding), colonies were fed daily at noon, and samples for RNA extraction were collected just prior to feeding. From four replicate colonies per condition, RNA was extracted separately from larval heads and the remaining body. Larval heads were isolated by cutting at the posterior prothoracic border with microdissection scissors. Each RNA sample consisted of a pool of ten individuals per tissue type per colony. The remaining replicate colonies per condition were allowed to complete their colony cycle to verify that IGEs on brood phenotypes occurred prior to conducting RNA-seq. From these colonies, larval developmental timing and survival was recorded, and all focal individuals were dissected and had their ovariole number recorded. We also measured the body size of 10 randomly selected individuals per colony.

### RNA extraction

Total RNA was extracted from fresh or flash-frozen tissue samples. Tissues were first placed in TRIzol (Thermo Fisher Scientific) on ice, then flash-frozen on dry ice and stored at −80 °C until use. After thawing samples on ice, tissues in TRIzol were homogenized using a TissueLyser II (QIAGEN) with stainless steel beads (2 min at 30 Hz). To enhance lysis efficiency, each sample then underwent a freeze–thaw cycle using dry ice and was homogenized a second time. Samples were then centrifuged at 21,300 × g for 1 min, and the lysate was transferred to Heavy Phase Lock Gel tubes (Quantabio), where chloroform (Sigma-Aldrich, C2432) was added for phase separation (0.2 mL chloroform per 1 mL TRIzol). Tubes were shaken for 15 s, incubated at room temperature for 2 min, and then centrifuged at 14,000 × g for 10 min at 4 °C. The aqueous, RNA-containing phase was transferred to a fresh tube and mixed with an equal volume of 70% ethanol. This mixture was then applied to a RNeasy Mini Kit column (QIAGEN) and processed according to the manufacturer’s protocol, including an on-column DNase digestion using RNase-Free DNase (QIAGEN) to remove residual genomic DNA. RNA concentration and integrity were assessed using an Agilent 2100 Bioanalyzer. Only samples with RNA integrity numbers (RIN) ≥ 7.0 were used for downstream applications (i.e., RNA-seq and RT-qPCR).

### RNA sequencing and quality control

RNA library preparation and sequencing were performed by the Genomics Resource Center at The Rockefeller University (RRID:SCR_020986). cDNA was synthesized from 1 ng total RNA using the SMARTer Stranded Total RNA-Seq Kit (Takara Bio) with rRNA depletion, and sequencing libraries were prepared using the Nextera XT DNA Library Preparation Kit (Illumina). Library fragment size distributions were assessed using an Agilent High Sensitivity D1000 ScreenTape Assay, showing an average insert size of ∼300 bp (excluding ∼120 bp adaptor). Libraries were multiplexed and sequenced in a single lane on an Illumina NovaSeq 6000 using an S2 flow cell to produce paired-end reads (2×100 bp).

Raw paired-end RNA-seq reads were quality filtered and adapter trimmed using fastp (*147*) (via the Rfastp wrapper). Reads were aligned to the *O. biroi* reference genome (*148*) (v5.4; GenBank assembly accession: GCA_003672135.1) using the subjunc function in the Rsubread package (*149*), and library and alignment quality metrics were evaluated using Picard (RnaSeqMetrics and AlignmentSummaryMetrics). Picard metrics were consistent with high-quality libraries, having predominantly exonic reads, low rRNA content, and high mapping rates. On average, 93.8% of paired-end reads passed quality filters and aligned to the reference genome per sample (range: 89.8–95.1%).

### Gene expression analyses

High-quality adapter-trimmed paired-end RNA-seq reads were quasi-mapped to the reference transcriptome using Salmon v1.10 (*150*) with GC bias correction enabled. Transcript-level abundance estimates generated by Salmon were aggregated to gene-level counts and TPM values using tximport (*151*), which mapped transcripts to genes based on the *O. biroi* annotation (GTF v5.4) (*152*). Principal component analysis (PCA) of variance-stabilized gene expression values, obtained using the vst function, was conducted to assess sample similarity and to detect potential outliers.

Gene-level counts were analyzed for differential expression using DESeq2 (*153*). Genes with low counts were excluded prior to differential expression analysis (genes required ≥10 counts in at least 12 of 16 samples). DESeq2 default dispersion estimation and independent filtering were applied. A generalized linear model (expression ∼ batch + caregiver genotype + larval genotype) was fitted to test for genotype effects while accounting for batch effects, where batch represented the day on which larvae were dissected and tissues were stored in TRIzol. RNA extraction for all samples from each tissue were performed on the same day, and all samples were pooled for RNA sequencing; therefore, no additional RNA extraction or sequencing batch variables were included. Significant differential expression was assessed using the Wald test with Benjamini–Hochberg false discovery rate (FDR) correction. Raw Salmon counts, R code used for differential gene expression analysis, and full annotated DESeq2 result tables are available at Zenodo.

### Gene set enrichment analysis and weighted gene co-expression network analysis

A curated Gene Ontology (GO) annotation was generated by aggregating gene-to-GO mappings from EggNOG, Ensembl, and UniProt databases. PFAM (Protein Families database) domain annotations were derived from InterProScan analysis of predicted proteins and mapped to gene identifiers for enrichment analyses.

Gene set enrichment analysis (GSEA) was conducted using the clusterProfiler (v4.18.4) R package (*154*). Using DESeq2 differential expression output, genes were ranked by their Wald statistics for enrichment testing of GO categories, and annotation tables were restricted to genes present in the ranked gene list prior to enrichment analysis. GO term definitions were obtained using the GO.db (v3.14.0) annotation package. PFAM domain enrichment was assessed using Fisher’s exact tests implemented with the enricher function of clusterProfiler. Benjamini–Hochberg corrections were applied to control the false discovery rate. GO enrichment maps were generated using the enrichplot (v.1.30.5) package.

Weighted gene co-expression network analysis (WGCNA) was performed with the WGCNA (v1.74) R package (*155*) using variance-stabilized gene expression values generated with DESeq2 from Salmon gene counts. Genes with low expression and variance were filtered prior to network construction, and the soft-thresholding power was determined using the WGCNA pickSoftThreshold function. Modules of co-expressed genes were identified using the blockwiseModules function. Module eigengenes were calculated and differential eigengene expression was assessed using limma (v3.66.0) (*156*), and PFAM domain enrichment tests were then conducted on genes belonging to modules of interest.

The curated genome GO and PFAM annotations as well as custom analysis scripts are available at Zenodo.

### RNA-FISH probe design

For *ilp1*, *ilp2*, *InR1* and *InR2*, RNA-FISH probes were custom designed as described by Glotzer et al. (*157*). All probe design code is available on GitHub (https://github.com/Social-Evolution-and-Behavior/Glotzer-Kronauer-2025), and custom probe sequences for our target genes are deposited in Zenodo. Oligonucleotide pools (50 pmol per oligonucleotide) were synthesized by IDT and dissolved in 50 μL nuclease-free water to produce a 1 μM stock solution.

For all other genes, probes were designed by Molecular Instruments. Probe lot numbers and HCR amplifiers are listed in Zenodo.

### RNA-FISH sample preparation and protocol

We performed the HCR v3.0 FISH protocol as described by Choi et al. (*158*), but with modifications and tissue preparation similar to that outlined in previous work (*152, 157*). Briefly, whole larvae were fixed in 4% paraformaldehyde in 1× PBS containing 0.1% Triton X-100 at 4 °C for 3 to 4 days. Dissected larval fat bodies were instead fixed for 2 hours at room temperature. After fixation, all samples were washed twice in 0.1% PBS-Tween for 10 minutes. To improve sample visibility and reagent permeability, whole larvae were then sonicated on a cooling block using a Q700 Sonicator (QSonica) fitted with a 1.6 mm microtip. Samples were sonicated at an amplitude of 10 with a pulse duration of 2 seconds on and 20 seconds off, for a total of 15 cycles. Samples then underwent three additional washes in 0.1% PBS-Tween for 10 minutes each, for a total of five washes.

Samples were pre-hybridized in HCR probe hybridization buffer (Molecular Instruments) at 37 °C for 30 minutes, followed by hybridization with probe mixtures in the same buffer for 48 hours at 37 °C with rotation (350 rpm) on a thermoblock. For each probe, 8 μL of 1 μM stock was added to 500 μL hybridization buffer. Up to four probes were multiplexed per sample, each paired with a distinct hairpin amplifier conjugated to fluorophores emitting at 488, 546, 594, or 647 nm. Post-hybridization, samples were washed five times for 10 minutes in preheated HCR probe wash buffer, followed by two 5-minute washes in 5× SSC with 0.1% Tween-20 (5× SSCT) at room temperature. Samples were then equilibrated in HCR amplification buffer for at least 10 minutes. Fluorescent hairpins were prepared by snap-cooling: 6 μL of each 3 μM hairpin stock was heated to 95 °C for 90 seconds, then cooled to room temperature in the dark for 30 minutes. Hairpins were combined in 100 μL amplification buffer and applied to samples. Amplification was performed for 16 hours at room temperature with rotation on a thermoblock (350 rpm) in the dark. Following amplification, samples were washed five times for 5 minutes each in 5× SSCT at room temperature and then incubated in DAPI (1:500 in 1× SSCT) for 8 hours. Samples were again washed five times for 5 minutes each in 5× SSCT at room temperature and mounted in SlowFade Glass (Thermo Fisher Scientific).

### Confocal microscopy

Tissues were imaged on a Zeiss LSM 900 confocal microscope equipped with 405 nm, 488 nm, 561 nm, and 633 nm laser lines, and operated with ZEN microscopy software. We used a Zeiss LD LCI Plan-Apochromat 40×/1.2 NA multi-immersion objective with glycerol immersion medium. Multi-channel Z-stacks of the tissues were acquired at 1.0× sampling density, ensuring Nyquist-optimal resolution. Optical sections were acquired at 0.25–0.75 μm intervals depending on tissue and sample, with a minimum resolution of 1024×1024 pixels. Detector gain, offset, and laser power were adjusted for each sample to optimize signal detection and prevent saturation. We used automated tile scanning in Zeiss ZEN with a minimum 10% tile overlap to capture images of larger samples. Tiles were stitched using the alignment algorithm implemented in ZEN, with the DAPI channel used for registration. Images were processed in Fiji (ImageJ) (*146*) for visualization only. Linear brightness and contrast adjustments were applied, followed by standard deviation Z-projections for each channel. Background subtraction was then performed prior to channel merging. Images are presented for qualitative localization, and no fluorescence intensity comparisons were made between samples. In each case, a single representative confocal image is shown, selected from at least three independent biological replicates.

To visualize expression in the larval body, we also stained for *obp1* and *csp3*, which showed relatively high levels of expression in the body based on our RNA-seq data (average of 28.5 and 763 TPM, respectively). Other than the fat body, we did not detect expression of *csp* or *obp* genes elsewhere in the larval body. However, because our RNA-FISH protocol does not label free floating hemolymph cells such as hemocytes, expression in these cells could not be assessed. Similarly, although we did not detect expression of examined neuropeptides or their receptors in other larval tissues, we cannot exclude the possibility that they are expressed elsewhere but were not detected under our experimental conditions. The original, unprocessed representative confocal image files are available at Zenodo.

### Artificial hemolymph preparation

Our functional experiments involved the preparation of *Drosophila* extracellular saline as described in Lu et al. (*159*), hereafter referred to as artificial hemolymph (AHL). The recipe is also available at Zenodo.

### Ecdysone feeding experiments

To investigate the effects of 20-hydroxyecdysone (20E) on larval growth and development, a stock solution was prepared by dissolving 5 mg 20E (Cayman Chemical, Cat# 16145) in 200 μL 100% ethanol (EtOH). For the working solution, 20μL of the 20E stock was mixed with 178 μL AHL and 2 μL blue food coloring (added for visual confirmation of ingestion), resulting in a final 20E concentration of 2.5 mg/mL. The control solution contained 20 μL 100% EtOH, while keeping AHL and dye volumes the same. Thus, both 20E treatment and control solutions contained 10% (v/v) ethanol. *Solenopsis invicta* brood was immersed in either the 20E or control solution overnight at room temperature, followed by an additional 24 hours at 4°C, and then stored at – 20°C until use.

For all iterations of this experiment, colonies were established with 10 random workers of unknown ages and 10 larvae. For genotype B (STC6) ants, colonies were fed treated food five times at 48-hour intervals, starting three days after larvae reached the fourth instar (*n* = 8 replicate colonies per treatment).

In experiments using genotype A (C16) ants, colonies were fed treated food three times at 48-hour intervals, starting three days after larvae reached the fourth instar, after which they were returned to a standard diet. The first experiment using genotype A ants included six replicate colonies per treatment (EtOH control vs. 20E). For the second iteration of the experiment using genotype A (C16), larvae from a subset of replicate colonies (*n* = 8 per treatment) were collected for quantitative gene expression analysis via RT-qPCR. Twenty-four hours after the third feeding, five larvae per colony were randomly selected and pooled for RNA extraction. The remaining colonies (*n* = 6 per treatment) continued their colony cycle so that developmental timing and final body size differences could be assessed.

### Corazonin injection experiments

To test the effects of Crz peptide on larval development, early fourth-instar larvae (within 48 hours of transitioning from the third to the fourth instar) of genotype A (C16) were microinjected with synthetic Crz peptide (Glp-TFQYSRGWTN-NH_2_; CPC Scientific, Cat# INSE-003A). The Crz injection solution consisted of 600 μM peptide, 0.5% DMSO, and 1% blue food coloring in AHL. The solvent-only control solution was identical in composition but lacked the peptide (i.e., 0.5% DMSO and 1% blue dye in AHL).

Injection solutions were loaded into injection needles pulled from quartz glass capillaries (Sutter Instrument, Cat# Q114-53-10NP). Injection needles were prepared using a P-2000 micropipette puller (Sutter Instrument), with the tips gently broken using microforceps. Injections were manually performed under a stereomicroscope using a FemtoJet 4i microinjector (Eppendorf) with 600-900 hPa injection pressure (p_i_) and 0 hPa compensation pressure (p_c_). Each larva received a ∼12.5 nL injection volume into their right dorsolateral prothoracic segment, delivering approximately 10 ng of Crz in treatment group individuals. The injection time for the desired volume was calculated for each needle by dispensing solution from the needle for 30 seconds onto parafilm and measuring the volume.

For each treatment condition (vehicle control and Crz), five replicate colonies were established with each colony containing 10 injected larvae, 5 young workers, and 10 older workers of unknown age. Larval developmental timing was monitored, and the body length and intercaste proportion of resulting adults was quantified. The experiment was then repeated, but larvae from a subset of replicate colonies (*n* = 8 per treatment) were harvested for gene expression analysis via RT-qPCR. For each RNA sample, five randomly selected larvae were collected from each replicate colony and pooled. The remaining replicate colonies were allowed to continue their colony cycle (*n* = 9 per treatment) for measurements of developmental timing and pupal body length.

### Quantifying relative gene expression using RT-qPCR

Primers were designed using Primer-BLAST (NCBI) to produce amplicons ranging from 71 to 186 bp, with target melting temperatures (Tm) between 55°C and 60°C. All qPCR primer sequences, as well as their amplification efficiencies and standard curve R^2^ values, are provided in Zenodo. All primers exhibited amplification efficiencies between 90–110%, with R^2^ above 0.98, indicating high-quality amplification across the dynamic range.

RNA quality and concentration of samples were assessed using an Agilent 2100 Bioanalyzer, and cDNA synthesis was performed using the SuperScript IV VILO Master Mix kit (Thermo Fisher Scientific; Cat# 11756050), with 400 ng of total RNA input per sample. Negative control reactions lacking reverse transcriptase (–RT controls) were included to confirm the absence of genomic DNA contamination. None of the primers tested amplified genomic DNA, and melting curve analyses confirmed a single Tm peak for each primer pair, indicating target specificity.

Quantitative PCR was performed in 10 µL reaction volumes using Applied Biosystems PowerTrack SYBR Green Master Mix (Thermo Fisher Scientific; Cat# 4309155) on a QuantStudio 5 Real-Time PCR System (Applied Biosystems). Reactions were assembled onto MicroAmp Optical 96-Well Reaction Plates (Cat #N8010560) and sealed with MicroAmp Optical Adhesive Film (Cat #4360954). All reactions were run in technical triplicate with no-template and –RT controls. Average Ct values were calculated from technical replicates following automatic outlier exclusion using default settings in QuantStudio 5 software. *GAPDH* served as the reference gene for normalization. Relative expression was calculated using the comparative Ct (ΔΔCt) method (*160*). Statistical analyses were performed on ΔΔCt values, and data were plotted as 2^(−ΔΔCt) to illustrate relative fold changes in gene expression. All Ct, ΔCt, ΔΔCt, and 2^(−ΔΔCt) values are provided in Zenodo.

### Statistical Analysis

Sample sizes, definitions of center and dispersion, and statistical tests employed are reported in the figure legends. Sample sizes are reported as number of colonies (*n*) and total number of individuals scored (*N*). Statistical significance was defined as *p* < 0.05. All models, including linear models (LMs), linear mixed models (LMMs), generalized linear models (GLMs), generalized linear mixed models (GLMMs), and cumulative link mixed models (CLMMs) were fit in R (v4.5.1) using the packages lme4 (v1.1.38) and ordinal (v2025.12.29) with colony identity treated as a random effect where applicable. Model fit was assessed using simulated residual diagnostics in the DHARMa (v0.4.7) package, including tests for overdispersion and overall goodness of fit based on residual uniformity.

Pupal size was analyzed using LMMs, with the nlme (v3.1.168) package used when necessary to model heterogeneous residual variance across conditions. Intercaste development was analyzed using binomial GLMMs. Developmental duration was analyzed using LMMs or CLMMs, depending on whether residual normality assumptions were met. Survival rates were analyzed using binomial GLMs, with beta-binomial GLMs employed using glmmTMB (v1.1.14) when overdispersion was detected. Unless otherwise noted, χ² values correspond to likelihood ratio tests (LRT), while F-statistics correspond to Type II ANOVAs conducted using the car (v3.1.5) package. Post hoc pairwise comparisons were conducted using estimated marginal means in the emmeans (v2.0.1) package, with p-values adjusted for multiple comparisons. Student’s t-tests, Welch’s t-tests, Mann–Whitney U tests, and two-way ANOVAs were conducted in GraphPad Prism (v10.5.0). For these tests, normality and homoscedasticity were assessed using GraphPad Prism’s residual diagnostics. For RT–qPCR analyses, two-tailed tests were used unless a directional effect was predicted based on prior evidence, in which case one-tailed tests were applied, and p-values were adjusted for multiple testing across 10 genes using the Benjamini–Hochberg FDR correction. All plots were generated either by GraphPad Prism (v10.5.0) or with R (v4.5.1) using the ggplot2 (v4.0.2) package. For box-and-whisker plots generated in R, center lines indicate the median, boxes indicate the interquartile range, and whiskers extend to 1.5 × the interquartile range (**Fig. 2B, 4B** and **fig. S1F**). For all remaining box and whisker plots, center lines indicate the median, boxes indicate the interquartile range, and whiskers extend to the minimum and maximum values.

Differential gene expression analyses performed using DESeq2 employed Wald tests, and p-values were adjusted using the Benjamini–Hochberg FDR method. Genes were considered differentially expressed at P_FDR_ ≤ 0.05 for larval body samples and P_FDR_ ≤ 0.1 for larval head samples. Quantile regression was used to compare median absolute log_2_ fold-change values between direct and indirect genetic effects. Linear models were used to evaluate associations between larval neuropeptide expression and *sad* expression, with caregiver genotype, larval genotype, their interaction, and *sad* expression included as fixed effects. Partial R^2^ values were calculated using rsq (v2.7) to quantify the variance uniquely explained by individual predictors. We applied Holm’s multiple comparisons correction when testing for differentially expressed WGCNA module eigengenes. For GO term GSEA and PFAM enrichment analyses, p-values were adjusted using the Benjamini–Hochberg FDR method.

## Supporting information

Supplementary Figures S1-S6; Supplementary Text; captions for Movies S1 & S2

## Acknowledgments

We thank Giacomo Glotzer for the custom design of RNA-FISH probes; Dominic Frank and Lindsey Lopes for training and assistance with confocal microscopy; Daniel H. Pastor for input on optimizing the RNA-FISH protocol; Kip Lacy for conducting genetic distance estimates between clonal genotypes; Yohann Chemtob for assistance with the behavioral tracking system; Alex Paul for advice on injections; and Orli Snir for advice on RNA extraction. We thank Taylor Hart for the cartoon ant illustrations. We are also grateful to Erich Jarvis, Li Zhao, and Vanessa Ruta for insightful feedback as members of P.K.P.’s thesis committee. We thank the Fisher Drug Discovery and Bioimaging Resource Center at Rockefeller University for access to equipment and training in its use. This is Clonal Raider Ant Project paper #47

## Author Contributions

P.K.P. and D.J.C.K. conceptualized the study and designed the experiments. P.K.P and S.V.-R. established and maintained experimental colonies. P.K.P and S.V.-R. collected body size, developmental timing, and survival data. J.L. contributed to data collection and analyses. P.K.P. developed and conducted the differential feeding assay and performed RNA extractions. P.K.P. and T.S.C. performed differential gene expression analyses, GSEA and WGCNA. P.K.P. and Y.C. conducted RNA-FISH stains. P.K.P. performed and analyzed the functional experiments and RT-qPCR. D.J.C.K. acquired funding and provided resources. P.K.P. and D.J.C.K. wrote the paper, and all authors read, edited, and approved the manuscript for publication. D.J.C.K. supervised the project.

## Data, code, and materials availability

RNA sequence data have been deposited to NCBI BioProject [Accession number: PRJNA1440417]. Fluorescence intensity measurements used to quantify food ingestion, RNA-seq gene counts, RT-qPCR data, and all body size, intercaste classification, developmental duration and survival data have been deposited at Zenodo and are publicly available as of the date of publication at [DOI: 10.5281/zenodo.18678443]. Unprocessed confocal microscopy images presented in this paper have been deposited at Zenodo and are publicly available as of the date of publication at [DOI: 10.5281/zenodo.18678443]. All original code has been deposited at Zenodo at [DOI: 10.5281/zenodo.18678443] and is publicly available as of the date of publication. Any additional information required to reanalyze the data reported in this paper is available from the lead contact upon request.

## Figures

Supplementary materials contain Figs. S1 to S6 and Movies S1 and S2.

