## Supplementary Figures S1-S6; Supplementary Text; captions for Movies S1 & S2 for "Caregiver genotype influences larval body size in ants by modulating ecdysteroid production"

**The Supplementary file includes:**

Supplementary Text

Figs. S1 to S6

Movies S1 to S2 (captions)

### Supplementary Text

Supplementary Note 1: Pyrokinin/PBAN receptor homology

The *O. biroi* genome (*1, 2*) contains two annotated receptors for PBAN-encoded peptides, both labeled pyrokinin-1 receptor (LOC105283619 and LOC105283116). To determine whether these receptors correspond to insect pyrokinin-1/diapause hormone receptors (PK1/DHr) or pyrokinin-2/PBAN receptors (PK2/PBANr) (*3*), we conducted BLASTP searches of each receptor against a selected panel of species (*Rhodnius prolixus*, *Tribolium castaneum*, *Aedes aegypti*, *D. melanogaster*, *B. mori*, *Apis mellifera*, *Monomorium pharaonis* and *O. biroi*) and used minimum-evolution distance–based clustering of sequence hits to assign receptor homology.

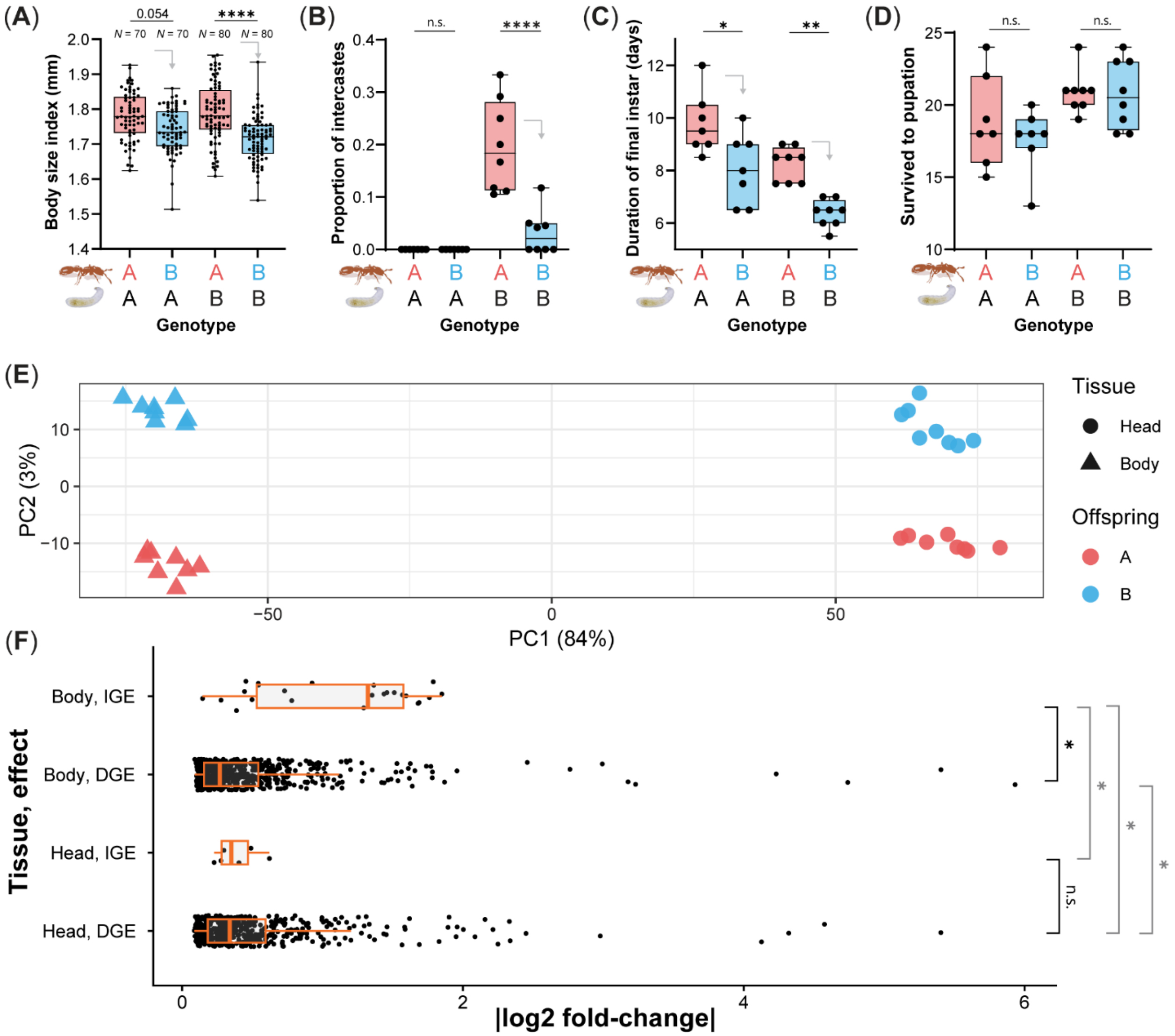

(A)
0.054
****
N = 70
N = 70
N = 80
N = 80
Body size index (mm)
2.0
1.9
1.8
1.7
1.6
1.5
1.4
A B A B
A A B B
Genotype
(B)
n.s.
****
Proportion of intercastes
0.4
0.3
0.2
0.1
0.0
A B A B
A A B B
Genotype
(C)
*
**
Duration of final instar (days)
12
10
8
6
A B A B
A A B B
Genotype
(D)
n.s.
n.s.
Survived to pupation
25
20
15
10
A B A B
A A B B
Genotype
(E)
PC2 (3%)
10
0
−10
−50
0
50
PC1 (84%)
Tissue
Head
Body
Offspring
A
B
(F)
Tissue, effect
Body, IGE
Body, DGE
Head, IGE
Head, DGE
*
*
*
*
n.s.
0
2
4
6
|log2 fold-change|

**Fig. S1. Direct and indirect genetic effects influence larval phenotypes and gene expression.** (**A-D**) Final body size, proportion of intercastes, duration of final instar, and survival of larvae across the four cross-foster conditions ($n$ = 7 to 8 colonies per condition). Only pairwise comparisons relevant to caregiver genotype effects are shown. On the x-axis, the color-coded letter in the top row indicates caregiver genotype, and the bottom row indicates larval genotype. (**A**) Body size was significantly affected by caregiver genotype (LMM: Type II Wald $\chi^2(1)$ = 31.12, $p < 0.0001$), but not by larval genotype ($\chi^2(1) = 0.44$, $p = 0.508$) or their interaction ($\chi^2(1) = 2.29$, $p = 0.130$). Points represent individual ants measured from multiple replicate colonies. (**B**) A two-way ANOVA on colony-level intercaste proportions showed a significant caregiver × larval genotype interaction ($F(1,26) = 20.23$, $p = 0.0001$). Genotype A caregivers coupled with genotype B larvae resulted in a significantly higher proportion of intercastes compared to all other conditions (AB vs AA, AB vs BA, AB vs BB; all Tukey-adjusted $p < 0.0001$). (**C**) A two-way ANOVA showed significant effects of both larval genotype ($F(1,26) = 21.11$, p < 0.0001) and caregiver genotype ($F(1,26) = 26.03$, p < 0.0001) on duration of the final instar, with no significant interaction effect ($F(1,26) = 0.05$, $p = 0.821$). (**D**) A two-way ANOVA showed a significant effect of larval genotype on larval survival ($F(1,26) = 9.04$, $p = 0.006$), but no significant effect of caregiver genotype ($F(1,26) = 0.67$, $p = 0.423$) or its interaction with larval genotype ($F(1,26) = 0.45$, $p = 0.508$). For (**B–D**), points represent individual colonies. (**E**) Principal component analysis (PCA) of DESeq2 variance-stabilized expression values calculated using all genes. Samples (total $n$ = 32) are separated by tissue type along PC1, with larval genotype segregating along PC2. Shapes indicate tissue type (head vs. body) and colors denote larval genotype (A vs. B). (**F**) Effect sizes (absolute $\log_2$ fold-change) of direct and indirect genetic effects (DGEs and IGEs, respectively) on genes showing differential expression ($P_{FDR}$ < 0.05) in the larval head or body. Quantile regression showed that, among differentially expressed genes, IGEs exhibited a significantly larger median effect size than DGEs in the larval body (median Δ = 1.02 in log2 fold change, Tukey-adjusted $p = 0.026$; in black). In contrast, no significant difference in median effect size was detected between IGEs and DGEs in larval head tissue (median Δ = 0.041, Tukey-adjusted $p = 0.987$; in black). Asterisks indicate significant differences based on Tukey-adjusted post hoc comparisons (*$p < 0.05$; **$p < 0.01$; ****$p$ < 0.0001).

(A)

| GeneID | Description (*Gene*) | log2FC | FC | *p* | *p. adj.* |
|---|---|---|---|---|---|
| LOC105284767 | **Cytochrome P450 315a1, Shadow (*sad*)** | -0.532 | 0.692 | 2.42E-09 | 2.59E-05 |
| LOC109610698 | Hypothetical protein | -0.747 | 0.596 | 4.69E-08 | 2.51E-04 |
| LOC105275477 | **PBAN-type neuropeptides (*pban*)** | -0.347 | 0.786 | 4.30E-06 | 0.015 |
| LOC105280204 | Uncharacterized, EF hand calcium binding domain 1 | -0.569 | 0.674 | 5.93E-06 | 0.016 |
| LOC105284706 | Spondin-1 (*Spon1*) | -0.288 | 0.819 | 9.58E-06 | 0.021 |
| LOC105275043 | 1-acyl-sn-glycerol-3-phosphate acyltransferase alpha (*AGPAT1*) | -0.445 | 0.734 | 1.96E-05 | 0.035 |
| LOC105279292 | Transmembrane protease serine 9 (*TMPRSS9*) | -0.609 | 0.656 | 3.38E-05 | 0.052 |
| LOC105280293 | **Pro-corazonin (*crz*)** | -0.478 | 0.718 | 4.72E-05 | 0.063 |
| LOC105279168 | **Prothoracicotropic hormone (*ptth*)** | -0.358 | 0.780 | 6.25E-05 | 0.074 |
| LOC105278524 | **Locust insulin-related peptide (*ilp2*)** | -0.289 | 0.819 | 0.002783 | 0.903 |
| LOC105287635 | **Juvenile hormone acid O-methyltransferase-like (*jhamt*)** | 0.305 | 1.236 | 0.005239 | 0.921 |

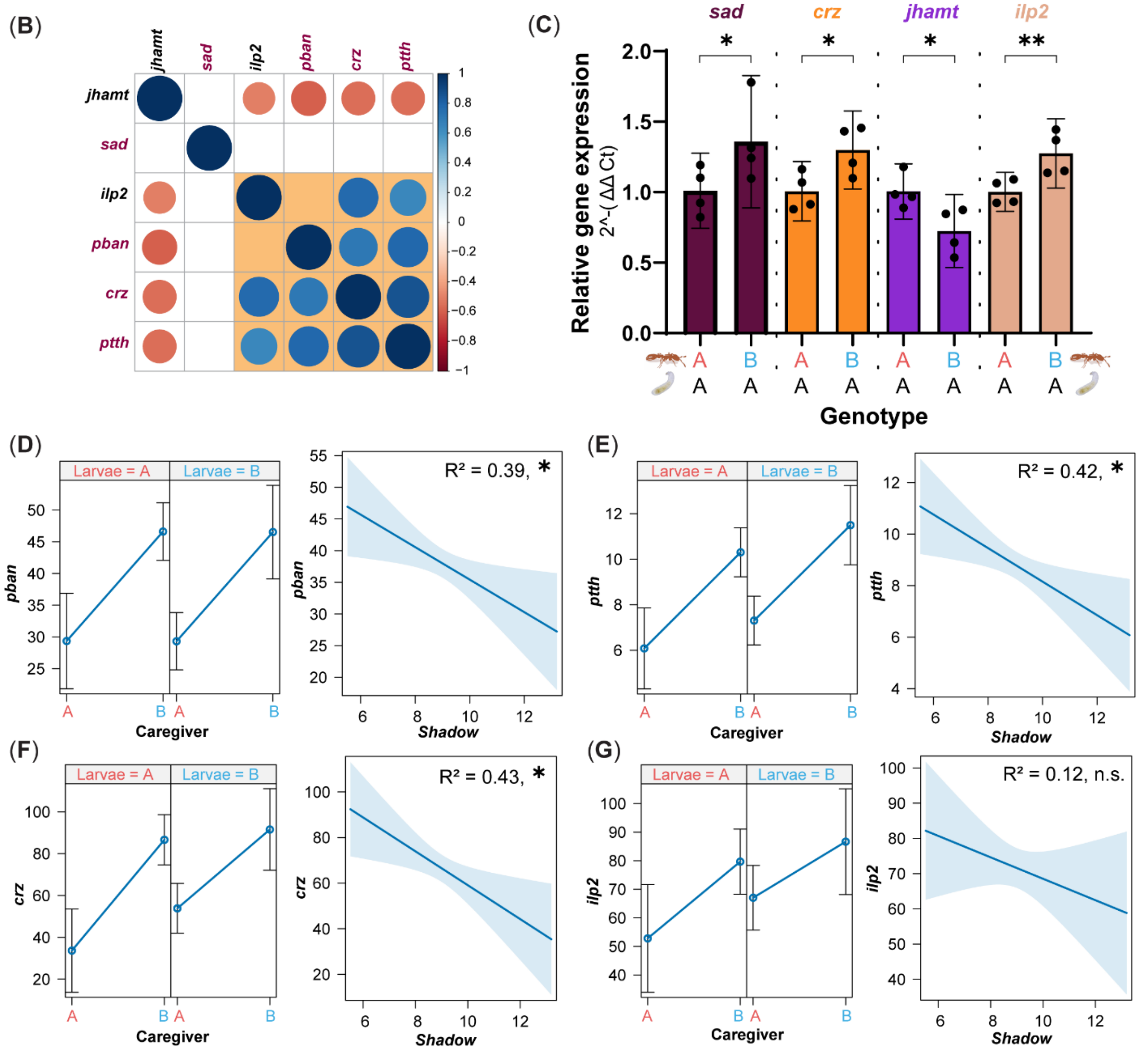

**Fig. S2. Correlated expression among neuropeptide and hormone biosynthesis genes affected by caregiver genotype in the larval head.**

(**A**) List of nine differentially expressed genes in the larval head due to caregiver genotype ($P_{FDR} < 0.1$), which includes multiple genes involved in regulating ecdysone production in insects (in bold purple text). Included are also *jhamt* and *ilp2*, two genes affecting growth in other insects. (**B**) Correlogram showing that *pban*, *ptth*, *crz* and *ilp2* expression are significantly correlated with each other (in orange shaded box), but not with *sad*. Both size and color of the circle represent the correlation coefficient, and only significant correlations ($p < 0.05$) are drawn. (**C**) Validation of bulk RNA-seq results using RT-qPCR, comparing the relative expression of *sad*, *crz*, *jhamt*, and *ilp2* for the same larval head RNA samples corresponding to genotype A larvae reared by caregivers of genotypes A or B ($n = 4$ per condition). All four genes were confirmed to be differentially expressed due to caregiver genotype (unpaired t-tests, one-tailed: *sad*, $t(6) = 2.19$, $p = 0.036$; *crz*, $t(6) = 2.69$, $p = 0.018$; *jhamt*, $t(6) = 2.61$, $p = 0.020$; and *ilp2*, $t(6) = 3.19$, $p = 0.009$). The color-coded letter in the top row indicates caregiver genotype, and the bottom row indicates larval genotype. Error bars represent 95% CIs of the mean. (**D–G**) Linear models (LMs) testing the effects of caregiver genotype, larval genotype, their interaction, and *sad* expression on larval neuropeptide expression. Left panels show estimated effects of caregiver genotype on neuropeptide expression for each larval genotype. Right panels show the relationship between neuropeptide expression and *sad* expression while controlling genotype effects. Bars and shaded areas represent 95% CIs of the mean. Partial $R^2$ values shown in each panel quantify variance in neuropeptide expression uniquely explained by *sad* after accounting for genotype effects. (**D**) Variation in *pban* expression was significantly explained by caregiver genotype (LM: $F(1,11) = 22.23$, $p < 0.001$) and *sad* expression ($F(1,11) = 6.99$, $p = 0.023$). (**E**) Variation in *ptth* expression was significantly explained by caregiver genotype (LM: $F(1,11) = 23.66$, $p < 0.001$) and *sad* expression ($F(1,11) = 8.04$, $p = 0.016$). (**F**) Variation in *crz* expression was significantly explained by caregiver genotype (LM: $F(1,11) = 21.54$, $p < 0.001$) and *sad* expression ($F(1,11) = 8.38$, $p = 0.015$). (**G**) Variation in *ilp2* expression was significantly explained by caregiver genotype (LM: $F(1,11) = 6.29$, $p = 0.029$), but not *sad* expression ($F(1,11) = 1.56$, $p = 0.238$).

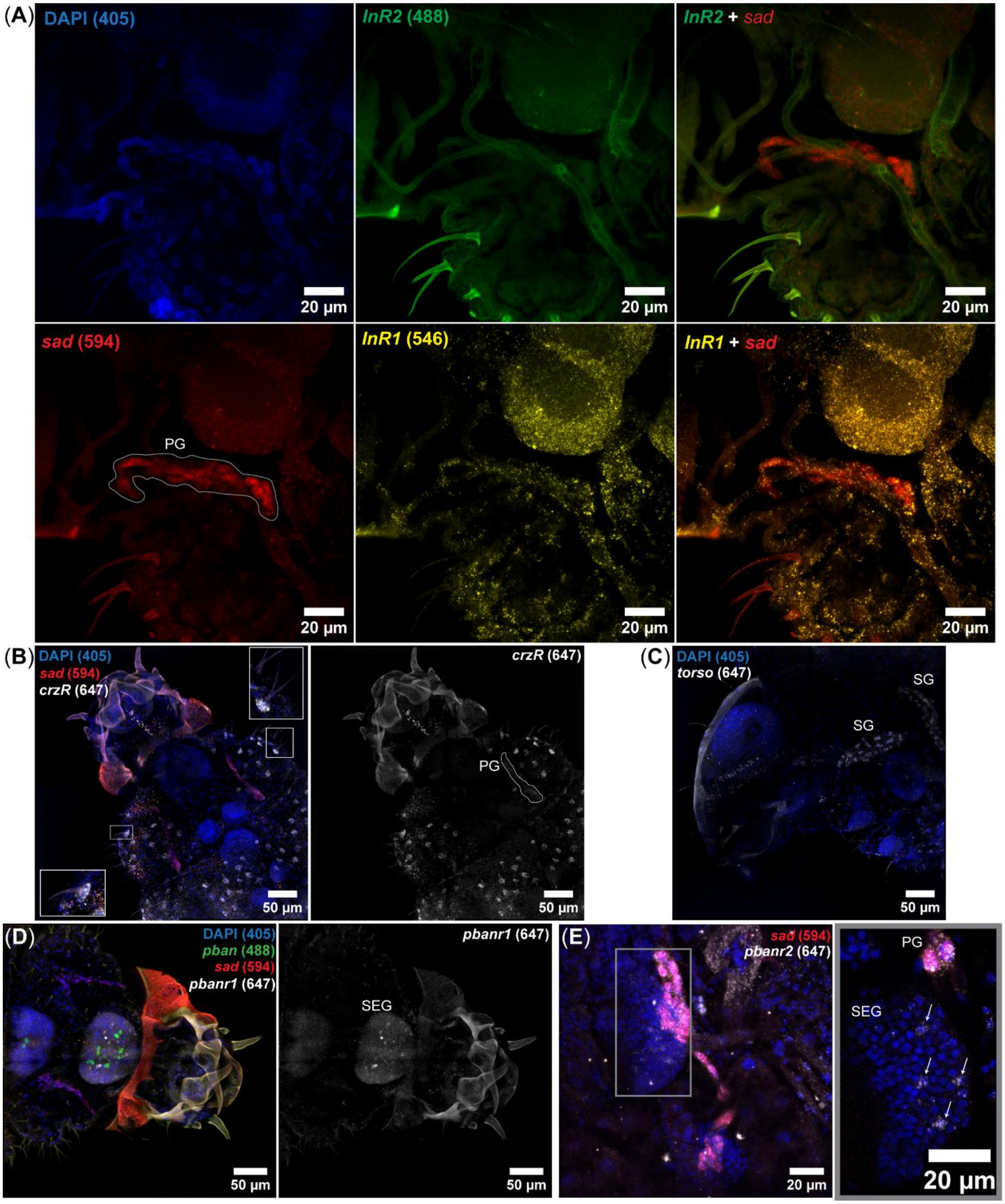
(A)
DAPI (405)
InR2 (488)
InR2 + sad
sad (594)
PG
InR1 (546)
InR1 + sad
20 µm
(B)
DAPI (405)
sad (594)
crzR (647)
PG
50 µm
(C)
DAPI (405)
torso (647)
SG
SG
50 µm
(D)
DAPI (405)
pban (488)
sad (594)
pbanr1 (647)
SEG
50 µm
(E)
sad (594)
pbanr2 (647)
PG
SEG
20 µm

**Fig. S3. *InR1*, but not *InR2*, is expressed by prothoracic gland cells.**

(**A**) Confocal image of an early fourth-instar larva co-stained for *sad*, *InR1* and *InR2* using RNA-FISH. Nuclei were counterstained using DAPI (blue; top left). The location of the prothoracic gland (PG) is marked by *sad* expression (red; bottom left). *InR2* expression was not detected in the larva (green; top middle), including the PG cells (merged *InR2* and *sad*; top right). *InR1*, on the other hand, was expressed ubiquitously (yellow; bottom middle), including in PG cells (merged *InR1* and *sad*; bottom right). (**B**) Confocal image of an early fourth-instar larva co-stained for *sad* and *crzR* (ventral view). In addition to PG cells, epidermal cells associated with setae express *crzR* (examples shown in inset boxes). Left, merged channels; right, *crzR* signal only (white). (**C**) Confocal profile view of a larval head showing *torso* expression in salivary gland (SG) cells. (**D**) Ventral view of a larva co-stained for *pban*, *sad*, and *pbanr1*. Multiple cells of the subesophageal ganglion (SEG) express *pbanr1*, and 16 SEG neurons express *pban*. Left, merged channels; right, *pbanr1* signal only (white). (**E**) Ventral view of the PG region co-stained for *sad* and *pbanr2*. In addition to expression in PG cells, *pbanr2* expression is detected in several SEG cells. On the right, a representative z-slice of the SEG corresponding to the grey boxed area is shown. Arrows indicate *pbanr2*-positive cells. Numbers in parentheses indicate laser excitation wavelengths. Receptor expression in additional tissues cannot be excluded.

(A)

| GeneID | Description (*Gene*) | | log2FC | FC | pvalue | padj |
|---|---|---|---|---|---|---|
| LOC105287745 | **odorant binding protein (*obp5*)** | ■ | 1.380 | 2.602 | 8.72E-29 | 9.04E-25 |
| LOC105286659 | **takeout-like (*jhbp*)** | ■ | 0.785 | 1.723 | 1.46E-19 | 7.56E-16 |
| LOC105287744 | **odorant binding protein (*obp6*)** | ■ | 1.375 | 2.594 | 1.04E-18 | 3.61E-15 |
| LOC105285832 | neuropeptide-like 3 | ■ | 1.606 | 3.044 | 6.94E-18 | 1.80E-14 |
| LOC105280989 | **ejaculatory bulb-specific protein 3 (*csp12*)** | ■ | 1.548 | 2.923 | 2.84E-16 | 5.90E-13 |
| LOC105288167 | pilosulin-1 | ■ | 1.927 | 3.802 | 3.52E-14 | 6.09E-11 |
| LOC109610747 | cuticle protein 18.7-like | ■ | 1.861 | 3.631 | 7.72E-14 | 1.14E-10 |
| LOC105284785 | odorant receptor co-receptor | ■ | 1.328 | 2.510 | 1.29E-11 | 1.68E-08 |
| LOC105280987 | **ejaculatory bulb-specific protein 3 (*csp14*)** | ■ | 1.486 | 2.802 | 2.42E-11 | 2.79E-08 |
| LOC105288165 | pilosulin-3a | ■ | 1.773 | 3.418 | 8.54E-10 | 8.86E-07 |
| LOC105286811 | uncharacterized, cytidine deaminase-like | | 1.870 | 3.654 | 1.16E-09 | 1.10E-06 |
| LOC105288166 | pilosulin-1-like | ■ | 1.682 | 3.208 | 3.81E-09 | 3.29E-06 |
| LOC105279397 | **odorant binding protein (*obp13*)** | ■ | 1.797 | 3.474 | 9.31E-09 | 7.43E-06 |
| LOC105285182 | larval cuticle protein A3A | ■ | 1.537 | 2.902 | 3.64E-08 | 2.70E-05 |
| LOC105278362 | **ejaculatory bulb-specific protein 3 (*csp1*)** | ■ | 0.502 | 1.416 | 4.82E-08 | 3.33E-05 |
| LOC105278344 | cytochrome P450 4g15 | ■ | 0.974 | 1.964 | 1.19E-06 | 7.73E-04 |
| LOC105277201 | sn1-specific diacylglycerol lipase beta | | -0.626 | 0.648 | 2.03E-05 | 0.012 |
| LOC105277445 | mRNA 3'-end-processing protein RNA14-like | | 1.066 | 2.093 | 2.82E-05 | 0.016 |
| LOC105277306 | forkhead box protein I2 | | -0.309 | 0.807 | 3.39E-05 | 0.018 |
| LOC105282332 | uncharacterized | | -0.816 | 0.568 | 5.09E-05 | 0.026 |
| LOC105277504 | mucin-2 | | 1.122 | 2.176 | 7.67E-05 | 0.038 |
| LOC113561855 | uncharacterized | ■ | 0.838 | 1.788 | 8.00E-05 | 0.038 |
| LOC105284725 | cytochrome P450 9e2 | | -0.719 | 0.608 | 1.06E-04 | 0.048 |
| LOC105278932 | venom allergen 3 | | 1.399 | 2.638 | 1.14E-04 | 0.049 |

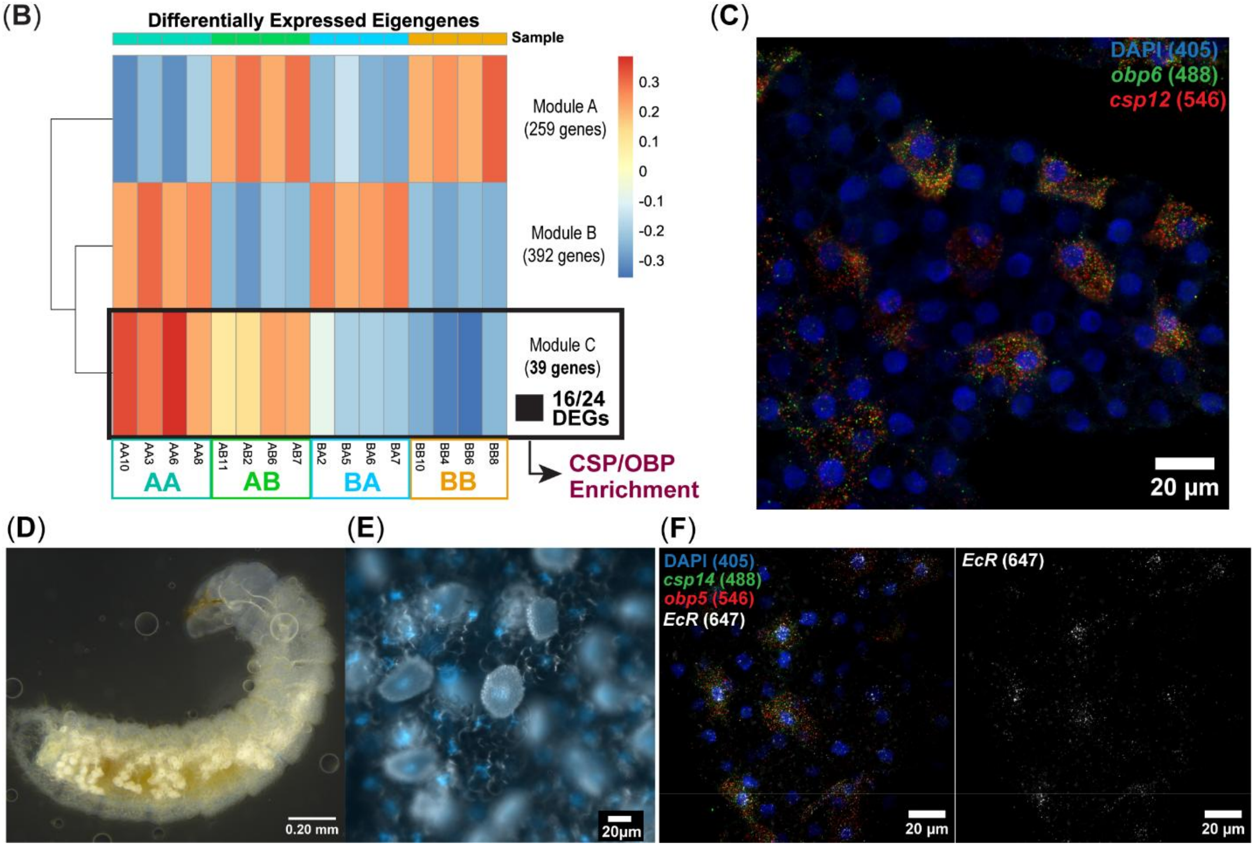

**Fig. S4. Lipid-binding carrier protein genes in the larval fat body exhibit correlated expression.**

(**A**) List of 24 genes differentially expressed in early fourth-instar larval bodies due to caregiver genotype, with secreted lipid-binding carrier proteins highlighted in bold purple text. Genes belonging to the protein families of CSPs and OBPs are significantly overrepresented (Pfam over-representation analysis, Fisher's test: *p.adj.* < 0.0001 for both). A positive $\log_2$ fold-change indicates relative upregulation in larvae reared by genotype A caregivers. (**B**) Three modules recovered from Weighted Gene Co-expression Network Analysis (WGCNA) show differential expression across conditions, with Module C representing a cluster of 39 genes that on average shows increased expression in larvae reared by caregivers of genotype A. This module is enriched for CSPs and OBPs (Fisher's exact test: CSPs, *p.adj* < 0.001; OBPs, *p.adj* < 0.0001), and it contains 16 of the 24 differentially expressed genes recovered by the DESeq2 analysis (denoted with black boxes in panel A). (**C**) RNA-FISH of *obp6* and *csp12*, showing enriched co-expression in a distinct population of cells in the larval fat body. (**D**) Brightfield stereomicroscope image of an *O. biroi* larva (lateral view). Conspicuous white spots are located throughout the larval body underneath the transparent cuticle. (**E**) Merged DAPI (blue) and transmitted-light DIC image of the larval fat body, showing that white matter is packed in nucleated cells. Transparent nucleated cells are trophocytes. (**F**) Larval fat bodies co-stained for *csp14*, *obp5*, and *EcR*. Left, composite image of all channels; right, *EcR* expression shown separately, enriched in the $csp^+/obp^+$ cells. Numbers in parentheses indicate laser excitation wavelengths.

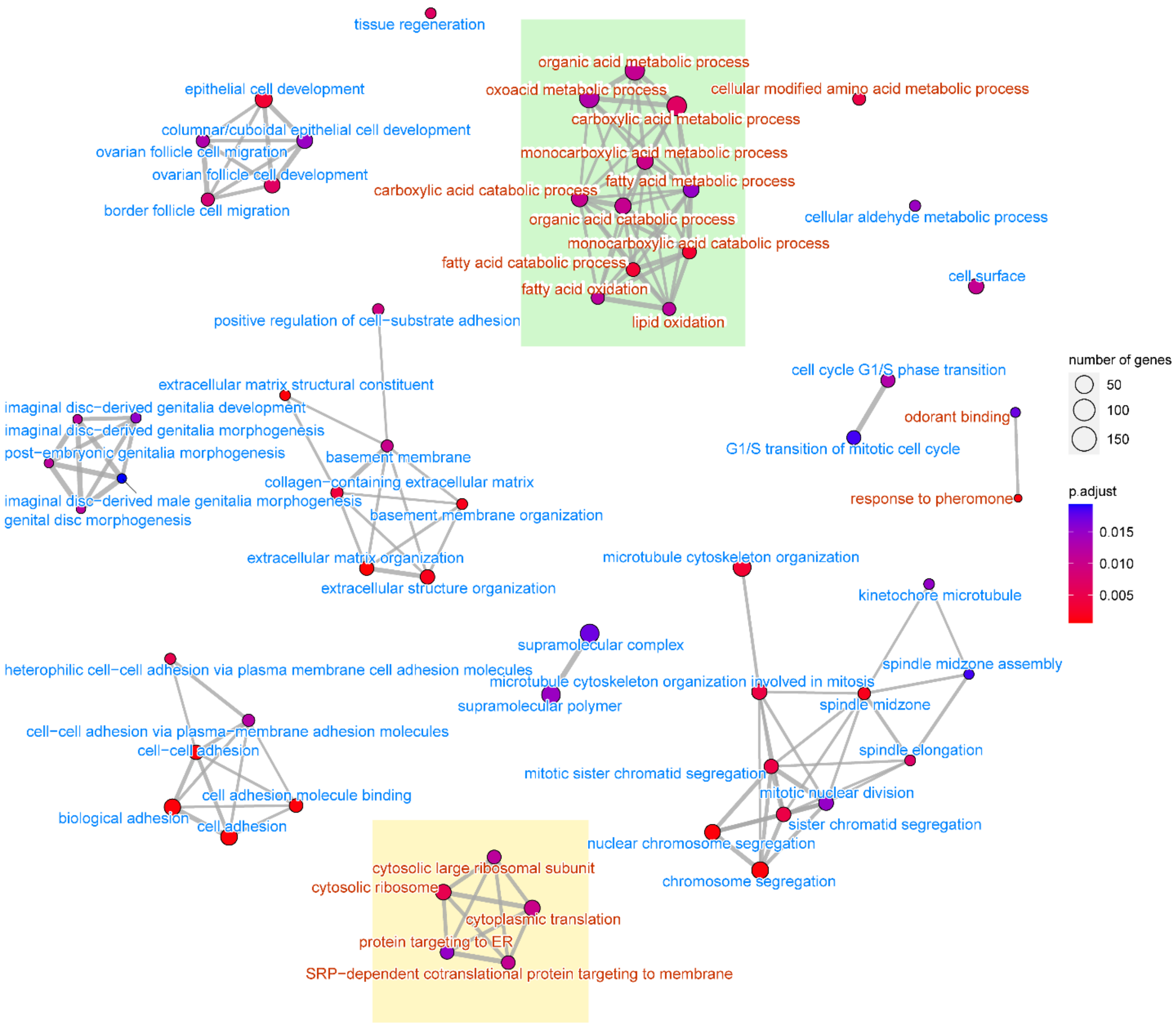


**Fig. S5. Caregiver genotype is associated with enrichment of fatty acid metabolism and secretory pathway gene sets in larvae**.

Enrichment map of the top 60 Gene Ontology (GO) terms from Gene Set Enrichment Analysis (GSEA) of the caregiver genotype effect in larval bodies. Nodes represent GO terms, with color indicating significance level and size corresponding to the number of associated genes. Edges indicate functional similarity based on shared genes, grouping related biological processes into clusters. Terms in red text are gene sets relatively upregulated in larvae reared by genotype A caregivers, and those in blue text are relatively upregulated in larvae reared by genotype B. The green and yellow boxes highlight clusters of GO terms associated with lipid catabolism and endoplasmic reticulum (ER)-directed translational activity, respectively. The translational activity cluster includes Signal Recognition Particle (SRP)-dependent cotranslational protein targeting and ER-directed translation, consistent with increased synthesis of secreted peptides. The leading-edge genes driving enrichment of monocarboxylic acid catabolic process include *HADH*, *ACADSB*, and *ACOX1*, which together support energy production via fatty acid oxidation.

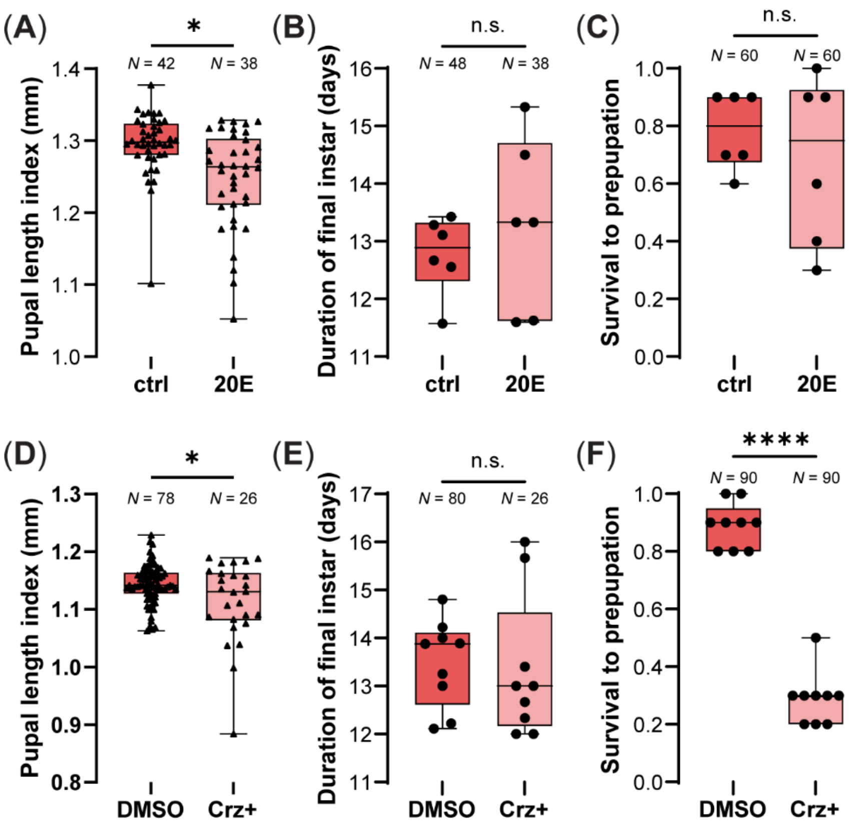


**Fig. S6. Ecdysone feeding and corazonin injection reduce body size without significant effects on developmental timing**.

(**A–C**) Effects of dietary 20E on (**A**) pupal size, (**B**) duration of final larval instar, and (**C**) larval survival ($n$ = 6 colonies per condition). Larvae were genotype A, and they were provided food supplemented with 20E a total of three times during the final instar. Dietary 20E resulted in smaller pupal sizes on average (LMM with treatment-specific residual variances: $\chi^2(1) = 5.74$, $p = 0.017$), with no detectable effects on duration of larval development (LMM: $\chi^2(1) = 0.71$, $p = 0.399$), or larval survival rates to prepupation (beta-binomial GLM: $\chi^2(1) = 0.25$, $p = 0.616$). (**D–F**) Phenotypic effects in the Crz injection experiment where larvae from a subset of colonies were collected for RT-qPCR analyses (see Fig. 6, F to H). Colonies ($n$ = 9 per condition) were established in parallel to quantify effects on (**D**) pupal body size, (**E**) duration of final instar, and (**F**) survival to prepupation. Crz injection reduced pupal body size relative to controls (LMM with treatment-specific residual variances: $\chi^2(1) = 3.95$, $p = 0.047$). Crz had no significant effect on final-instar duration (CLMM: $\chi^2(1) = 0.04$, $p = 0.844$) but significantly decreased larval survival (binomial GLM: $\chi^2(1) = 72.82$, $p < 0.0001$). Triangles correspond to individual ants, while circles correspond to replicate colonies. Asterisks indicate significant differences (*$p < 0.05$; ****$p < 0.0001$).

**Movie S1. Dorsal view of *ilp1* and *ilp2* expression in the larval head.**

Head of fourth-instar larvae, co-stained for *ptth* (yellow), *ilp2* (red), and *ilp1* (white) using RNA-FISH confocal microscopy. Nuclei were counterstained using DAPI (blue). *Ptth* was expressed in two bilateral pairs of dorsolateral neurons. *Ilp1* was bilaterally expressed in a population of cells located in the anterodorsal region of the brain. In contrast, *ilp2* was expressed in a bilateral cluster of cells located medially, corresponding to the canonical insulin-producing cells.

**Movie S2. Ventral view of *ilp1* and *ilp2* expression in the larval head.**

Head of fourth-instar larvae, co-stained for *ptth* (yellow), *ilp2* (red), and *ilp1* (white) using RNA-FISH confocal microscopy. Nuclei were counterstained using DAPI (blue). *Ptth* was expressed in two bilateral pairs of dorsolateral neurons. *Ilp1* was expressed in several disparate cells in the brain ventral to the medial insulin-producing cells expressing *ilp2*.

### Supplementary Information References

1. P. R. Oxley *et al.*, The Genome of the Clonal Raider Ant *Cerapachys biroi*. *Curr. Biol.* **24**, 451-458 (2014).
2. S. K. McKenzie, D. J. C. Kronauer, The genomic architecture and molecular evolution of ant odorant receptors. *Genome Res.* **28**, 1757-1765 (2018).
3. X. Dou, R. Jurenka, Pheromone biosynthesis activating neuropeptide family in insects: a review. *Front. Endocrinol.* **14**, 1274750 (2023).
